# Zebrahub – Multimodal Zebrafish Developmental Atlas Reveals the State-Transition Dynamics of Late-Vertebrate Pluripotent Axial Progenitors

**DOI:** 10.1101/2023.03.06.531398

**Authors:** Merlin Lange, Alejandro Granados, Shruthi VijayKumar, Jordão Bragantini, Sarah Ancheta, Sreejith Santhosh, Michael Borja, Hirofumi Kobayashi, Erin McGeever, Ahmet Can Solak, Bin Yang, Xiang Zhao, Yang Liu, Angela M. Detweiler, Sheryl Paul, Honey Mekonen, Tiger Lao, Rachel Banks, Yang-Joon Kim, Adrian Jacobo, Keir Balla, Kyle Awayan, Samuel D’Souza, Robert Haase, Alexandre Dizeux, Olivier Pourquie, Rafael Gómez-Sjöberg, Greg Huber, Mattia Serra, Norma Neff, Angela Oliveira Pisco, Loïc A. Royer

## Abstract

Elucidating the developmental processes of organisms requires a comprehensive understanding of cellular lineages in the spatial, temporal, and molecular domains. In this study, we introduce Zebrahub, a dynamic atlas of zebrafish embryonic development that integrates single-cell sequencing time course data with lineage reconstructions facilitated by light-sheet microscopy. This atlas offers high-resolution and in-depth molecular insights into zebrafish development, achieved through the sequencing of individual embryos across ten developmental stages, complemented by trajectory reconstructions. Zebrahub also incorporates an interactive tool to navigate the complex cellular flows and lineages derived from light-sheet microscopy data, enabling *in silico* fate mapping experiments. To demonstrate the versatility of our multi-modal resource, we utilize Zebrahub to provide fresh insights into the pluripotency of Neuro-Mesodermal Progenitors (NMPs). Our publicly accessible web-based platform, Zebrahub, is a foundational resource for studying developmental processes at both transcriptional and spatiotemporal levels, providing researchers with an integrated approach to exploring and analyzing the complexities of cellular lineages during zebrafish embryogenesis.

## INTRODUCTION

The development of a fertilized egg into a multicellular organism requires the coordination of various processes, including genetic regulation, cellular movements, and tissue morphodynamics. A major challenge for developmental biology has been understanding the lineages of individual cells as they organize at various scales to form tissues and organs. The investigation of cell lineages in embryos started over a century ago using light microscopes to track cells visually^1, 2^. A key milestone was the publication in 1983 of the complete cell lineage of *C. elegans*^3, 4^. However, the reconstruction of entire lineage trees is still limited to organisms with highly stereotypic cell fates and good optical access^5^. Advances in fluorescence light-sheet microscopy have enabled *in-toto* imaging of embryonic development at cellular resolution using zebrafish, mice, and drosophila, allowing for the reconstruction of specific lineages based on cell positions over time^6–10^. Single-cell RNA-sequencing (scRNAseq) technologies have provided complementary means of tracking cell lineages based on their transcriptomes over developmental time^11–21^. Here, we combine light-sheet microscopy with scRNAseq of individual developing zebrafish embryos to construct a comprehensive multimodal atlas of cell lineages and molecular states at unique spatiotemporal resolution. We provide the data and analytical tools as an integrated and publicly available resource: Zebrahub. To validate the quality and applicability of Zebrahub’s datasets, we focused our investigation on Neuro-Mesodermal Progenitors (NMPs). NMPs constitute a crucial and evolutionarily conserved bipotent population instrumental in axial elongation. These progenitors yield skeletal muscles and posterior neural cells, yet their developmental dynamics remain ambiguous^22–27^. Through Zebrahub, we uncovered indications of a state transition in NMPs’ fate dynamics at both transcriptional and live-imaging tissue levels.

## RESULTS

### Time-resolved single-embryo deep scRNAseq dataset of zebrafish development

Zebrahub’s sequencing resource consists of a dataset containing 120,444 cells from 40 individual zebrafish embryos and larvae spanning ten developmental stages from end-of-gastrulation embryos to 10-day larvae (10-, 12-, 14-, 16-, 19-, 24-hours post-fertilization, as well as 2-, 3-, 5- and 10-days post-fertilization). To capture single-cell RNA-sequencing variability across individuals, the cell-dissociation protocol was optimized to avoid embryo pooling, thus achieving non-pooled single-embryo single-cell RNA sequencing (Fig. 1a, Supp. Fig. 1 & Methods). Four fish were sequenced for each of the ten developmental stages, thus obtaining 40 individually sequenced embryos. Using strict quality-control criteria (Supp. Fig. 2 and Methods), we got a least one thousand high-quality single-cell transcriptomic profiles for the early developmental time points and more than 20 thousand cells at later stages (Fig. 1a – left). Importantly, we obtained a minimum of 3,000 genes per cell (25,000 UMIs per cell, see Supp. Fig. 2) for developmental stages before 24 hpf – highlighting the depth of our dataset.

**Figure 1.**
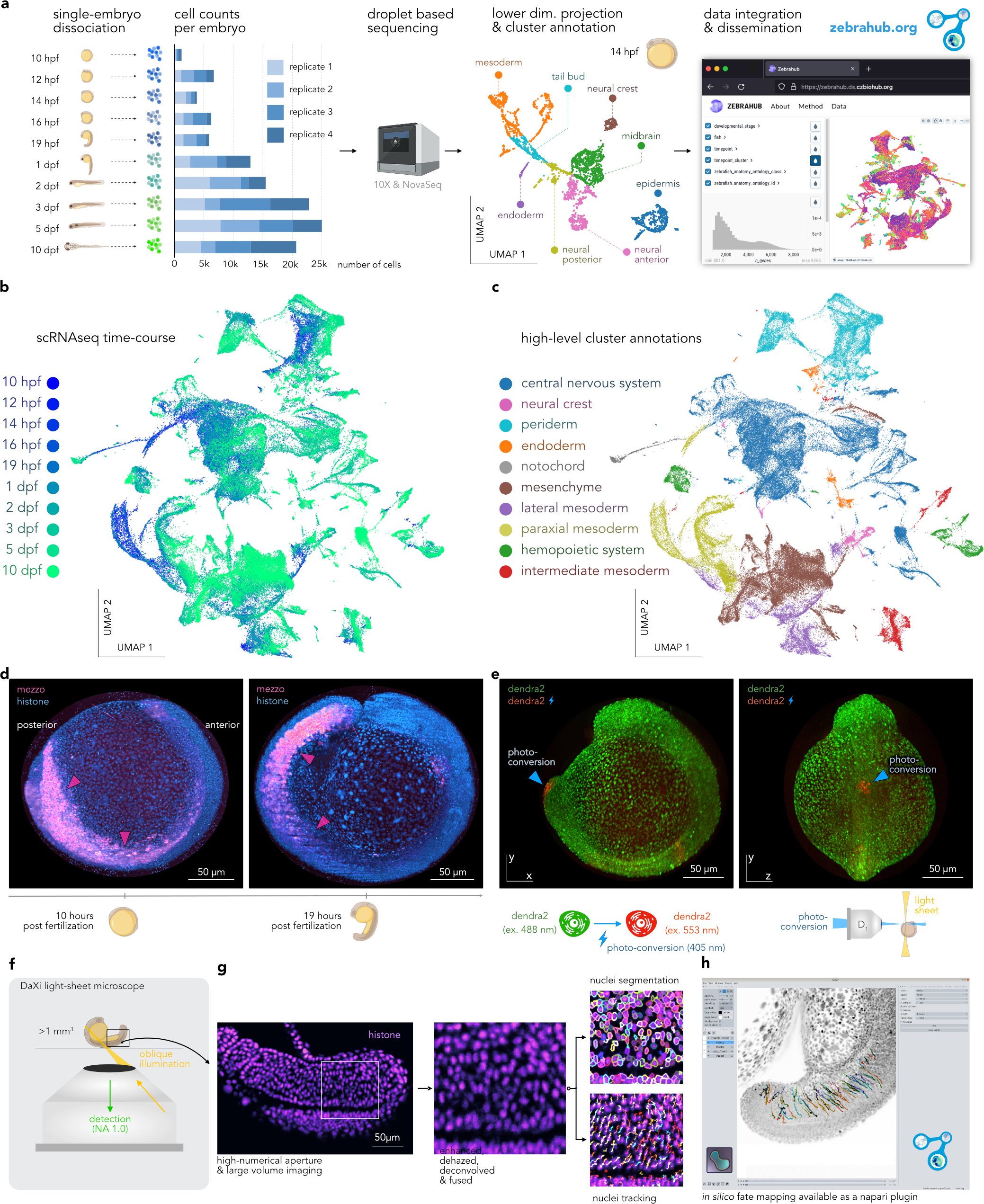
Zebrahub – Multimodal Zebrafish Developmental Atlas. **(a)** Zebrahub’s scRNAseq experimental pipeline for single-zebrafish-embryo single-cell RNA sequencing across developmental stages (dpf: days post fertilization, hpf: hours post fertilization). From left to right, we show: First, the number of cells per time point and replicates. Second, an example UMAP for a 14 hpf embryo transcriptome colored by cell type (3862 cells total). Third, Zebrahub’s web portal allows the exploration of scRNAseq data online. **(b)** UMAP representation of the entire dataset of 40 single-embryo transcriptomes colored by developmental stage (120k total cells). **(c)** Same but colored by annotated cell types. **(d)** Maximum intensity projections of a 12-hour light-sheet live-imaging of a zebrafish embryo expressing *histone*-mCherry (cyan) and *mezzo*-GFP (magenta). Arrows indicate the *mezzo*-positive cells. **(e)** Similar imaging but on an embryo expressing the photoconvertible protein Dendra2. Photoconverted cells are in red (blue arrows). **(f)** Overview of DaXi – a high-resolution, large field-of-view and multi-view single-objective light-sheet microscope. **(g)** Computational pipeline for image processing and nuclei tracking applied to images of a histone-mCherry zebrafish embryo acquired with our DaXi microscope (see Methods). **(h)** Screenshot of the *in silico* fate-mapping *napari* plugin available as part of Zebrahub (Methods and Supp. Fig. 15).

**Figure 2.**
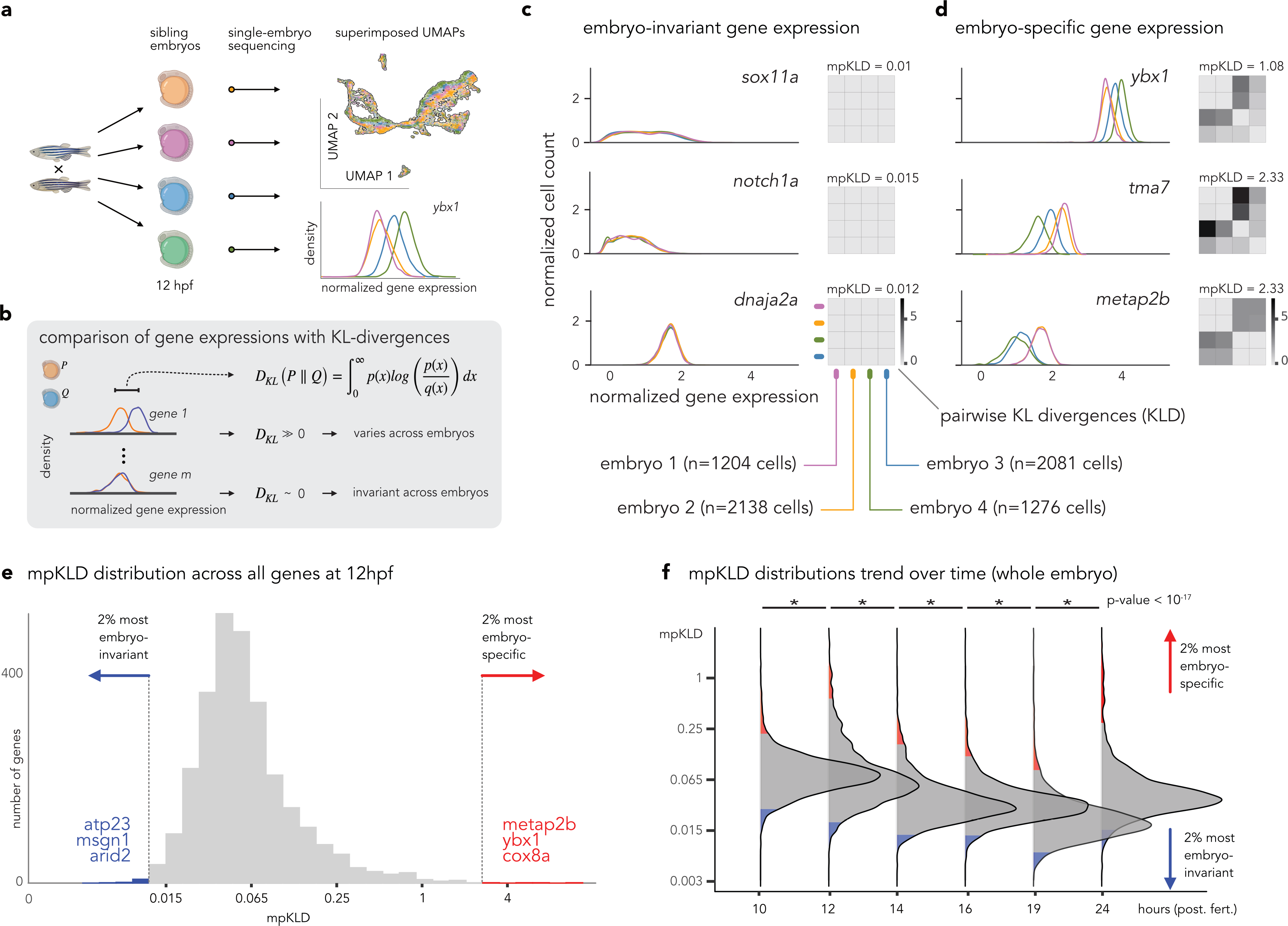
Inter-individual transcriptomic variability analysis. **(a)** Sequencing of sibling embryos enables the study of inter-individual gene expression variance. A UMAP of the single-cell data without batch correction (12 hpf, four embryos, 6699 cells total). Example of normalized expression of gene *ybx1* for four embryos (color coded). **(b)** We used the Kullback-Leibler divergence (KLD) to quantify the pairwise similarity between probability distributions, illustrating variant (top, KLD ≫ 0) and invariant (bottom, KLD ≈ 0) gene expression across embryos. **(c)** Example distributions for embryo-invariant genes (KLD ≈ 0) at 12 hpf: *sox11a*, *notch1a*, and *dnaja2a*. The x-axis indicates the log-normalized expression values, and the y-axis shows the density (relative number of cells). Heatmaps show all pairwise KLD values. We summarized these heatmaps per gene by the median pairwise KLD (mpKLD, values above the heatmap). **(d)** Example distributions for embryo-variant genes (KLD ≫ 0) at 12 hpf: *ybx1*, *tma7*, *metap2b*. **(e)** Histograms of mpKLD values scores for all cells at 12 hpf. A few examples of high and low mpKLD genes are shown. **(f)** Distributions of mpKLD values across all genes at 10, 12, 14, 16, 19, and 24 hpf. In **e** and **f,** the top and bottom 2% of genes are indicated as colored areas (p-values < 10^−15^ are marked with an asterisk).

### Cell-type clustering and annotation

To visualize the transcriptomes for these 120,444 cells, we generated uniform manifold projections (UMAP^28^) for each developmental stage (Fig. 1a – center and Supp. Fig. 3) and the whole dataset (Fig. 1b). Branching patterns that recapitulate the progression of embryonic development were observed, with cells taken from early embryos located at the beginning of branches and cells obtained from older juveniles positioned towards the outer areas (Fig. 1b and Supp. Video 1). A total of 529 cell clusters (Leiden clustering) were annotated by examining enriched genes and cross-referencing with the published literature and the zebrafish database ZFIN^29^ (Supp. Table 1). The identified cell types were then grouped into higher-level tissue types (Fig. 1c and Supp. Video 2). The resulting projections for the different developmental stages reveal an increase in the number of cell types over time consistent with cell differentiation during development (Supp. Fig. 4). The entire dataset, including projections and annotations, is available for download and online browsing through a user-friendly portal at zebrahub.org (Fig. 1a – right).

**Figure 3.**
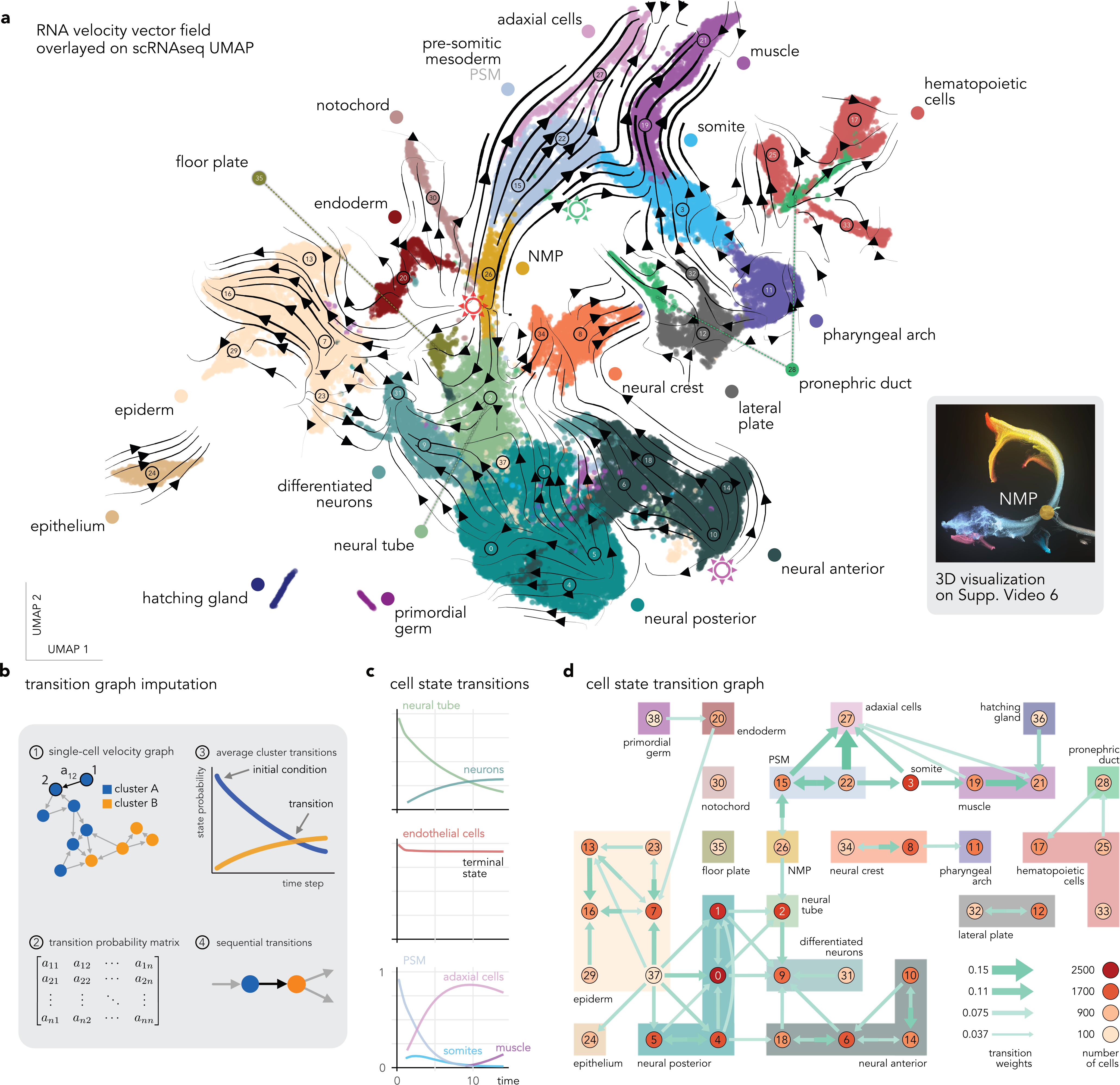
Whole embryo RNA velocity and derived cell state transition map. **(a)** Projection (UMAP) of zebrafish embryo transcriptomes (early development, 10 to 24 hpf time points), color-coded by cell type. Arrow streams correspond to the average RNA velocities of single cells. **(b)** Workflow to reconstruct state transitions from RNA velocity graph. _①_ Weighted directed graph represents the expected transitions between transcriptional states from single cells. _②_ This graph defines a Markov process that can be represented as a matrix. _③_ We can then estimate cluster transitions by simulating successive state transitions for groups of cells through the graph. _④_ Finally, we summarize cluster-level transitions as a coarse-grained graph. **(c)** Examples of average transition probabilities for neural tube, endothelial cells, and mesodermal tissue. **(d)** A coarse-grain graph of cell state transitions shows transitions between all cell states. The width of the arrows is proportional to the transition rates. Colors and numbers correspond to the annotation in **a**. Nodes grouped by a bounding box indicate clusters of cells that belong to the same broad cell types; colors correspond to those in **a**.

**Figure 4.**
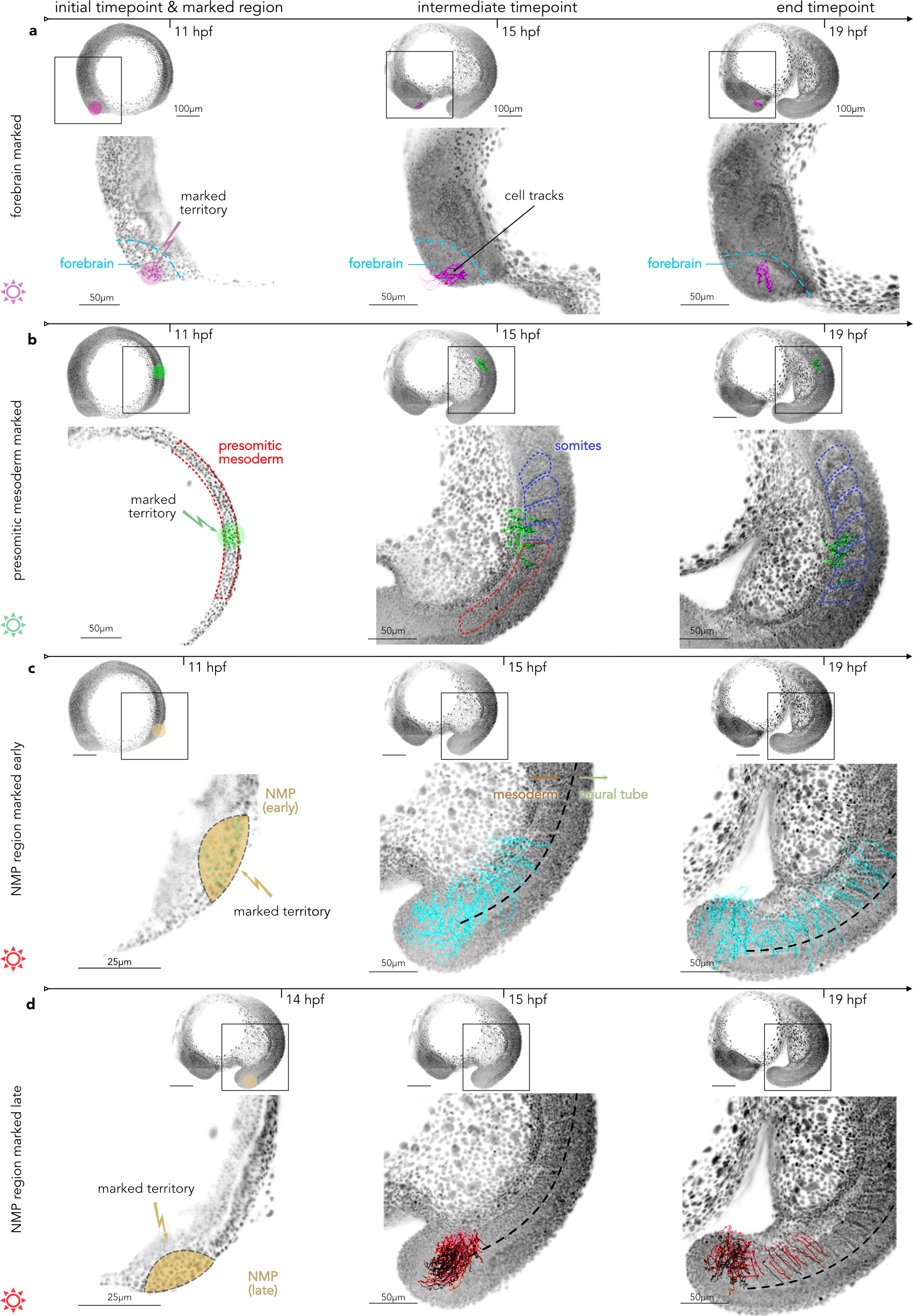
Reconstruction of the zebrafish embryonic spatiotemporal lineages via *in silico* fate mapping. **(a)** *In silico* fate mapping of a population of forebrain cells. Left - selected territory at the initial timepoint, middle - intermediate timepoint showing tracks for selected cells, right – final time point. The purple star * refers to the RNA velocity in Fig. 3a. **(b)** *In silico* fate mapping of the posterior mesodermal progenitors in the presomitic mesoderm. We observe that these cells integrate into the somites that will then generate muscles. **(c)** *In silico* fate mapping of the pluripotent early NMP territory. **(d)** *In silico* fate mapping of the late presumptive NMP territory. These cells primarily integrate the mesodermal tissue (70% - Supp. Fig. 14 e).

### Single-cell image-based fate mapping during zebrafish development

To complement the sequence-derived transcriptomic state information, the individual cellular dynamics in developing embryos were imaged with an adaptive multi-view light-sheet microscope^30, 31^ (Supp. Fig. 5a). To accelerate the dissemination of this technology, both the hardware design and control software are open-source (Supp. Fig. 5b to g, Methods). This platform lets us image the spatiotemporal cellular dynamics among cells of interest using transgenic reporter lines, as illustrated using a pan-cellular marker (transgenic line *mezzo*:eGFP^32^, see Fig. 1d and Supp. Video 3). However, transgenic reporter lines for all lineages of interest are not readily available nor easily engineered. To follow groups of cells and tissues of interest in the absence of reporter lines, our instrument can also label specific subsets of cells by photo-converting fluorophores such as Dendra2 (Fig. 1e).

**Figure 5.**
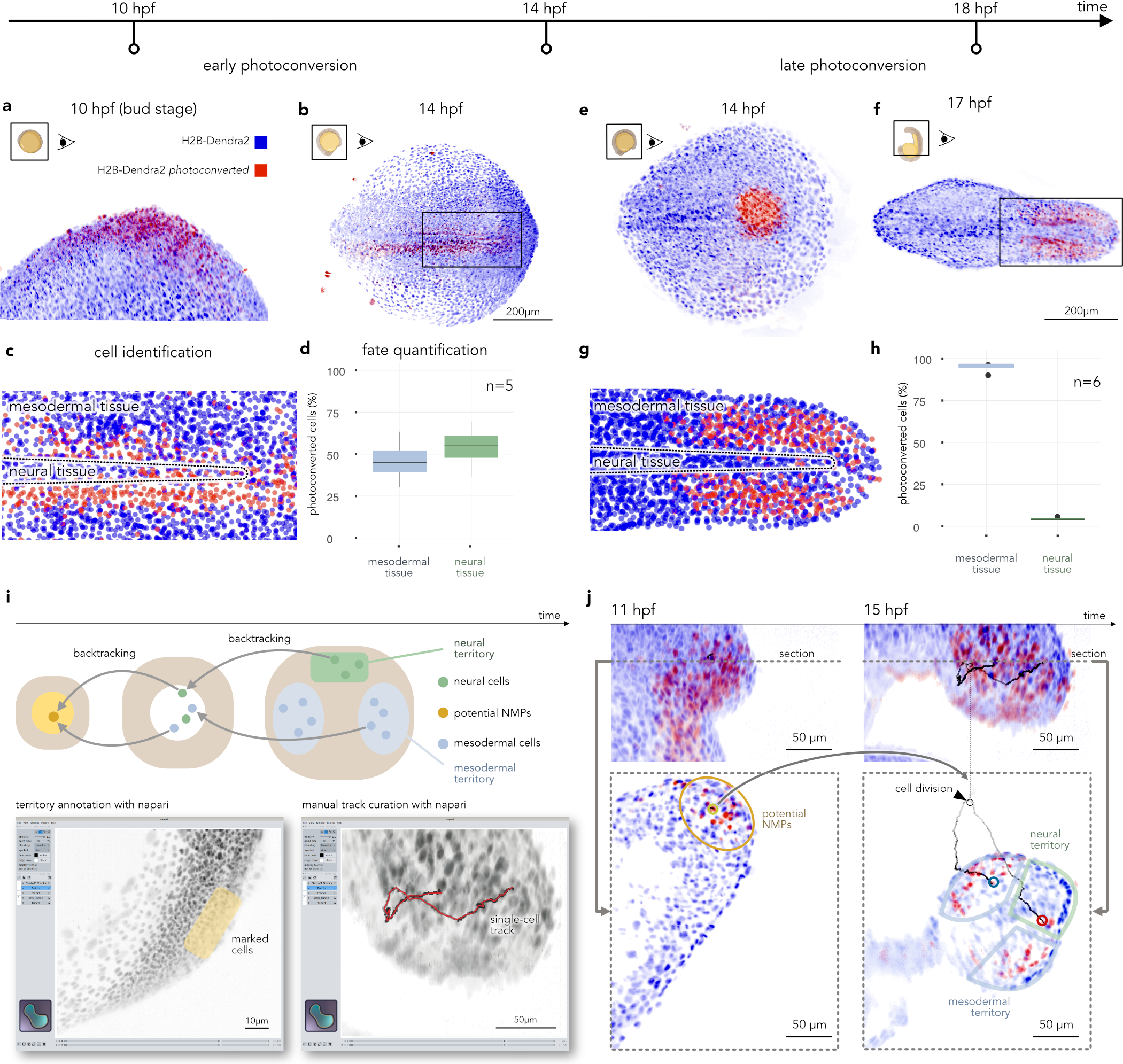
*In vivo* fate mapping of NMPs via photo-conversion and live imaging. **(a)** Photoconversion of the early presumptive NMP territory at ten hpf. **(b)** Distribution of the photoconverted cells and their progeny at 14 hpf. **(c)** Segmentation and identification of the photoconverted cells in the area selected in **b** (rectangle). **(d)** Distribution of early photoconverted cells and their progeny with mesodermal (left) and neural (right) fates (n= 5 animals). **(e)** Photoconversion of the late NMP territory at 14 hpf. **(f)** Distribution of the photoconverted cells and their progeny cells at 17 hpf. **(g)** Segmentation and identification of the photoconverted cells in the area selected in **f** (rectangle). **(h)** Distribution of the late photoconverted cells and their progeny (n=6 animals). **(i)** Identification of NMPs from territory annotations and backtracking inferred from automatically generated tracking data (top panel). Screenshot of the *napari* plugin for *in silico* fate mapping, showing territory annotation (left) and 3d cell track curation (right). **(j)** Illustration of a single early NMP (left) dividing into two daughter cells. After cell division, the first daughter cell integrates the neural territory, while the second integrates into the mesodermal territory (right). The top panels are two maximal-intensity projections of the posterior zebrafish embryo. The bottom panels are section visualization.

### High-resolution light-sheet imaging for lineage reconstruction

To image and accurately track the movement of cells in densely populated and rapidly developing embryos, we employed a single-objective light-sheet microscope (DaXi) capable of high-resolution imaging (1.0 NA) over large volumes (> 1 mm^3^) without the need for time-consuming sample preparation protocols^33^ (Fig. 1f, Supp. Fig. 6a, b). An ubiquitous histone reporter line driving the expression of mCherry^34^ was used to image the nuclei of cells. Following their acquisition, images were processed and enhanced to facilitate the identification and tracking of cell nuclei (Fig. 1g and Supp. Video 4). Our cell-tracking pipeline uses a neural network to detect cell nuclei in the images and delineate boundaries between cells. This information is then used to obtain nuclei segmentations and the corresponding tracks (Fig. 1g, Supp. Video 4 and methods). An *in silico* fate mapping plugin was developed for the multidimensional image viewer *napari*^35^ to help users explore and use these extensive datasets (Fig. 1h and Supp. Fig. 12). This software allows users to conduct virtual fate mapping experiments by marking territories and following the trajectories of cells and their descendants. All imaging datasets and tools are accessible online at zebrahub.org and, together with the scRNAseq datasets, constitute a foundational multimodal resource to study vertebrate development.

**Figure 6.**
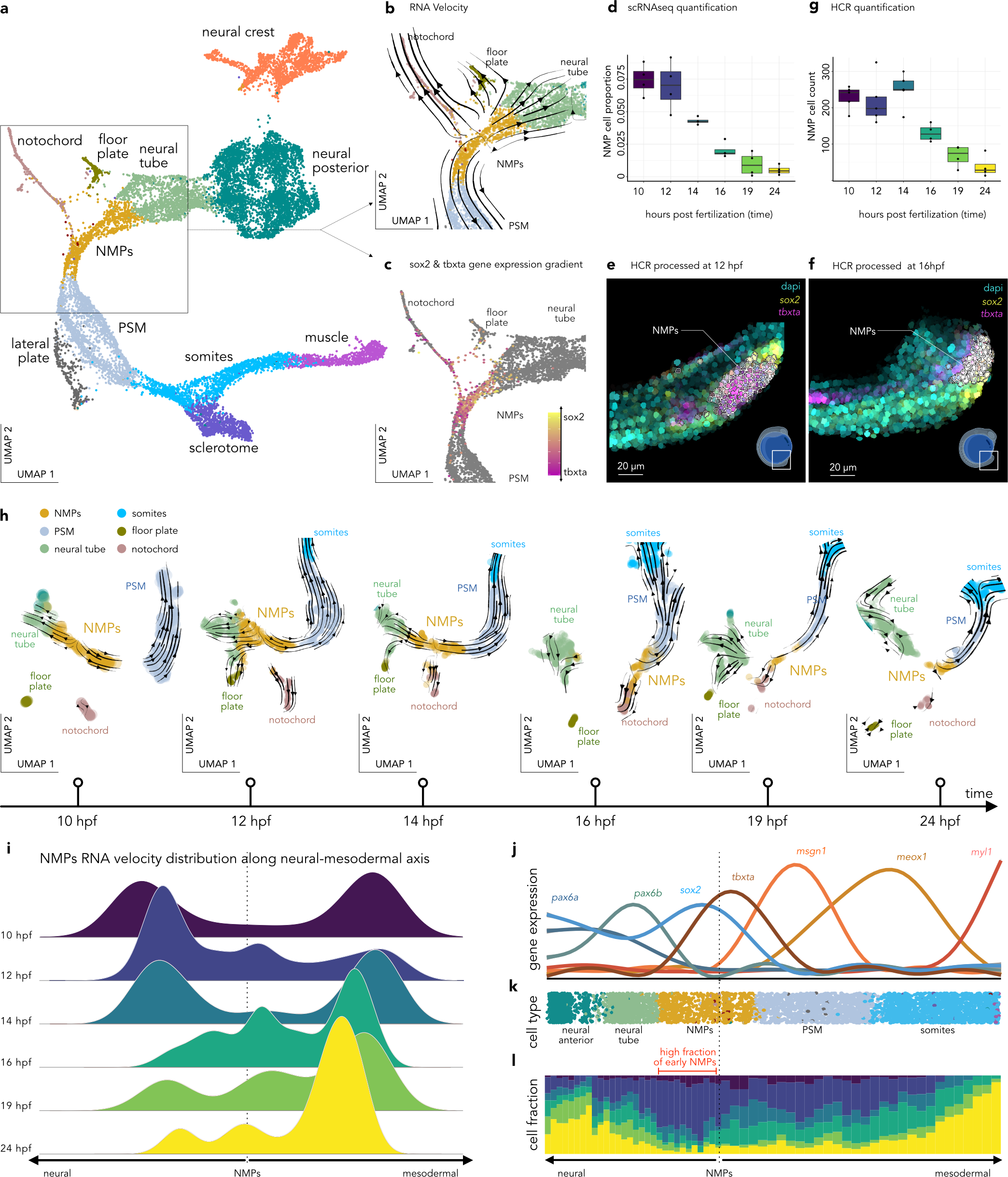
Transcriptional dynamics of NMPs during early development. **(a)** Reprojection (UMAP) of the NMPs and adjacent clusters, color-coded by annotated cell types. **(b)** Zoom in on the NMPs with color-coded cell types and superimposed RNA velocity streamlines. **(c)** Same projection but with cells color-coded by their *sox2/tbxta* expression ratio; stronger yellow indicates higher *sox2* and stronger magenta indicates higher *tbxta*. **(d) Quantifying** the proportion of cells in the NMP clusters relative to the total number of cells for each developmental stage. **(e)** Expression of *sox2* (yellow) and *tbxta* (red) for segmented cells (12 hpf) derived by hybridization chain reaction (HCR). Cells co-expressing *sox2* and *tbxta*, presumptive NMPs, have their boundary in white. **(f)** Same as **e** but at 16 hpf. **(g)** Quantification of presumptive NMPs (cells co-expressing *sox2* and *tbxta*) obtained from the HCR image analysis. **(h)** Velocity trajectories are projected into a time series of aligned UMAPs centered around the NMPs and their adjacent clusters, colored by cell type. **(i)** Time-resolved quantification of the distribution of RNA velocity for NMPs along the neural-mesodermal axis. These are the normalized distributions of the magnitudes of individual RNA velocity vectors pointing toward either the neural or mesodermal tissues. **(j)** Pseudotime resolved scaled expression of genes involved in neural development (left axis) and mesodermal development (right axis). **(k)** Annotated cell-type visualized on the pseudotime axis rooted in the NMPs, recapitulating the neural (left) and the mesodermal (right) developmental transcriptomic trajectories. **(l)** Cell fraction from different developmental stages visualized along the pseudotime axis.

### Inter-individual transcriptomic variance between sibling embryos

The single-embryo resolution and high sequencing depth of the scRNAseq datasets (Supp. Fig. 2) provide a unique opportunity to study the variability of gene expression during development by comparing the single-cell transcriptomes of siblings (Fig. 2a). To quantify the differences in gene expression patterns between siblings, a framework comparing gene expression distributions across cells was developed, utilizing the Kullback-Leibler divergence^36^ (Fig. 2b). A Kullback-Leibler divergence of 0 indicates that the two distributions are identical, while values greater than zero indicate increasing divergences between the distributions (Fig.2b). For example, at 12 hpf, multiple genes showed seemingly identical expression patterns between siblings (*e.g., sox11a*, *notch1a* and *dnaia2a* in Fig. 2c), whereas other genes showed embryo-specific distributions (*e.g., ybx1*, *tma7* and *meta2b* in Fig. 2d). We computed all pairwise Kullback-Leibler divergences for each gene and summarized these using the median value (mpKLD for median pairwise Kullback-Leibler divergence, Fig. 2c and Fig. 2d gray heatmaps - Methods). At 12 hpf, most genes have very similar expression levels across embryos (mpKLD≈0.06), with a few genes exhibiting extremely low mpKLDs (<0.01) – indicating nearly identical gene expression across embryos (bottom 2% of genes colored in blue in Fig. 2e). However, a tail of genes with high mpKLDs (>2) was also observed – indicating highly varying gene expression across siblings (top 2% of genes colored in red on Fig. 2e). Applying the same method, genes with identical or embryo specific expression were observed at different time points and in particular tissues (Supp. Fig. 7). We confirmed that the mpKLD between siblings could not be explained by sampling error by showing that they are significantly different from those obtained from a null distribution (see Methods and Supp. Fig. 8a). Comparable mpKLD distributions were observed for the other early stages (Supp. Fig. 8b). These results indicate that using single embryo resolution enables the analysis of transcriptomic variability. Interestingly, we observed that most genes exhibit similar expression levels across embryos of the same age.

**Figure 7.**
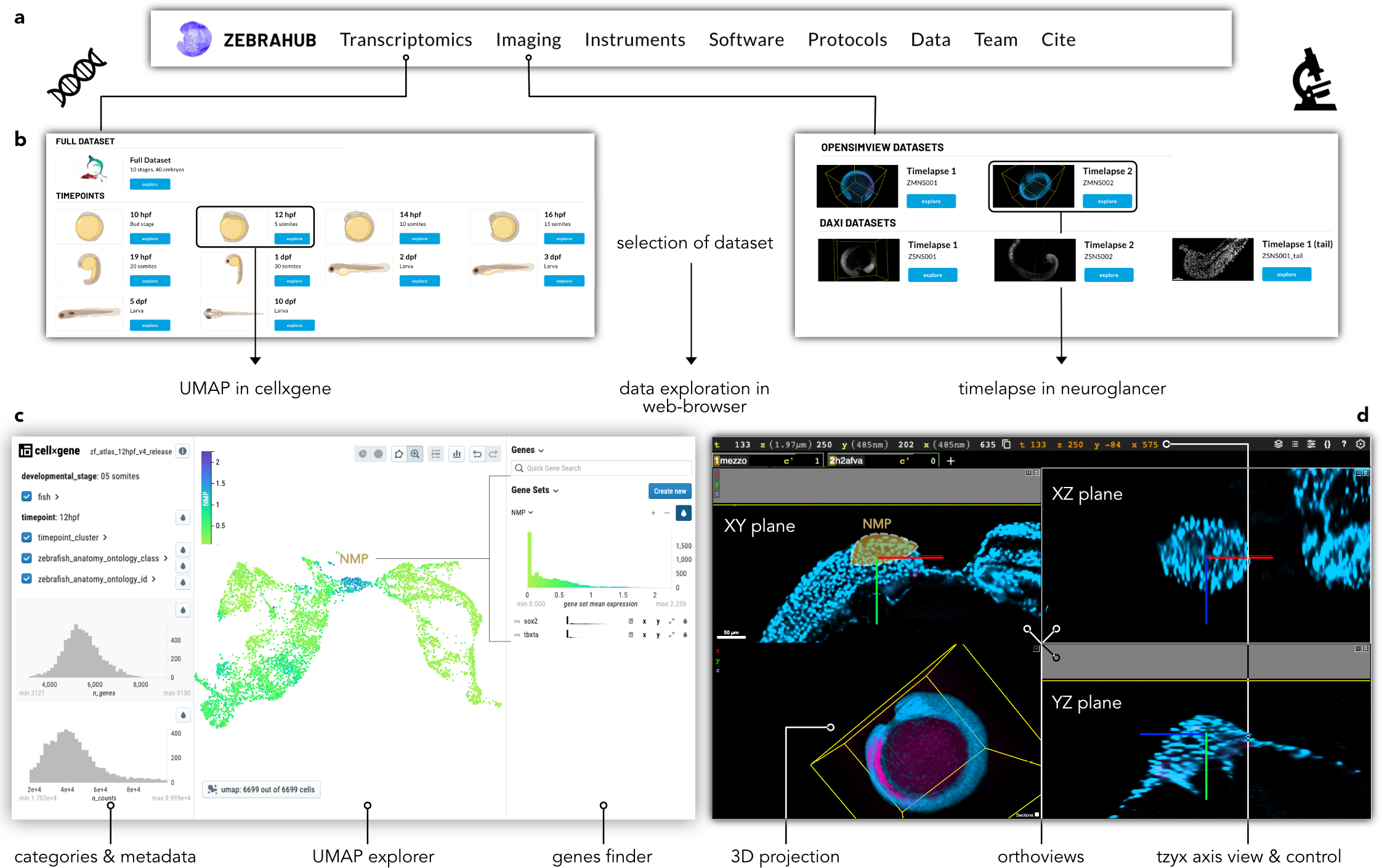
The Zebrahub website. **(a)** Zebrahub main tabs: Explore various datasets, instruments, software, protocols, and raw data. **(b)** Examples of in-depth exploration of Zebrahub high-dimensional datasets: scRNAseq (left) and light-sheet time-lapse (right). **(c)** Left: Zebrahub 12 hpf scRNAseq Atlas on the *cellxgene* platform, to visualize metadata, explore the annotated UMAP, search, and create gene sets (colored example: NMPs expressing sox2 and tbxta). Right: multiview light-sheet time-lapse exploration using *neuroglancer*. highlight the importance of image-based cell lineage reconstructions to complement transcriptomic studies to validate cell lineages. Zebrahub’s unique combination of datasets and technologies was instrumental in refining our understanding of lineages and thus helping answer finer questions on the factors influencing cellular fate. Specifically, Zebrahub enabled us to characterize the transition dynamics of axis elongation progenitors precisely. We proposed a model of the fate dynamics of NMPs that incorporates data from transcriptomic states, image-derived lineages, and tissue kinematics. We discovered that during axis elongation, NMPs transition from contributing to multiple mesodermal and neural lineages to exclusively contributing to the mesoderm. A peak expression of key molecular pathway’s components (Wnt, FGF, and RA pathway) and the presence of the pluripotency factor oct4, linked to the maintenance and differentiation of axial progenitors^50, 57–59^ were found in early pluripotent NMPs. Combining in silico and in vivo image-based fate mapping confirmed the fate restriction from pluripotent to exclusively mesodermal progenitors at around 15 hpf. Altogether, these results challenge the traditional view of lineage commitment post-gastrulation and do support the concept of late and broad differentiation plasticity.

### Time-resolved analysis of transcriptomic variability between individuals shows that gene expression reaches the highest level of similarity among siblings at 19 hpf

To better understand transcriptomic variability, the analysis was extended to six other early development time points (10 to 24 hpf). The mpKLD distribution peaks for all time points are low (between 0.065 and 0.015), indicating that most genes showed similar expression across embryos. Interestingly, we observed an overall decrease in mpKLD, as shown by a distribution shift (Figure 2f – Two-Sample Kolmogorov-Smirnov Test across consecutive time points, p-value < 10^−15^). The gene expression distributions across siblings reach a point of minimal gene expression variability at 19 hpf, with a rebound in variability at 24 hpf (Fig. 2f). This pattern was also observed when examining specific cell types (Supp. Fig. 9a). We speculate that the higher degree of similarity at 19 hpf reflects the tight control of gene expression at the end of the segmentation period^38^.

### Zebrahub RNA velocity recapitulates lineage formation

To gain a deeper understanding of gene expression dynamics during early developmental stages (10 hpf to 24 hpf), RNA velocity^39^ was computed. The RNA velocity vectors were projected on the annotated UMAP (Fig.3a and Supp. Video 5) and recapitulated the progression of embryonic time (Supp. Fig. 10a). For example, the RNA velocities clearly show the mesodermal differentiation trajectory, starting from pre-somitic mesoderm, through somites, to muscles (Fig. 3a, upper right, green star *). Interestingly, the cluster at the origin of the mesodermal branch, indicated by a red star *, is also at the origin of a branch going to the neural tissue. The cells composing this cluster are enriched for both neural and mesodermal transcription factors (*sox2* and *tbxta*, respectively, see Supp. Fig. 10b, c). These transcription factors are the molecular signature of the Neuro-Mesodermal Progenitors (NMPs) – a recently characterized population of late pluripotent axial progenitors that are believed to contribute to mesoderm and neural tissue post-gastrulation^22, 40^.

### Cell state transition graph analysis reveals transcriptomic cellular dynamics during embryogenesis

To substantiate these findings quantitatively and avoid confounding effects due to the UMAP projection, we coarse-grained our RNA velocity data into a cell state transition graph representing the dynamics of differentiation between cell types and tissues (Fig. 3b). Modeling the single-cell transcriptome transitions from the RNA velocities as a Markov process allowed us to explore a variety of cell type trajectories^41^ (see Methods and Fig. 3c). Cell states related to progenitor populations transitioned into their corresponding differentiated states (e.g., neural tube to neurons in Fig. 3c – top). In contrast, matured or fully differentiated cell states did not transition into any other state (e.g., endothelial cells in Fig. 3c – middle). The cell state transitions were recapitulated in a coarse-grain graph (Fig. 3d). Interestingly, the cell transition graph confirms well-known lineages such as the mesodermal lineage (PSM ↘ somites ↘ muscle). The graph also supports the notion that NMPs transition to both mesodermal and neural states (Fig. 3D, red star *). Intriguingly, the graph also reveals bidirectional transitions between cell types that are proximal in transcriptional space but are not known to be part of the same developmental lineages (e.g., primordial germ and endoderm; pronephric duct and hematopoietic system, Fig. 3D). Overall, our cell state transition graph recapitulates the main developmental lineages. It also possibly suggests novel lateral transitions between early cell types that do not correspond to known spatiotemporal developmental lineages.

### Image-based developmental lineage reconstruction using *in silico* fate mapping

RNA velocity provides valuable information about the direction of cell state dynamics in transcriptomic space. However, as described earlier, it is essential to note that predictive transcriptomics does not always align with developmental lineages. Therefore, it becomes necessary to employ systematic image-based lineage reconstruction to fully capture the cellular spatiotemporal dynamics during development. A machine-learning-based algorithm for nuclei segmentation and tracking was used to reconstruct high-resolution fate maps from single-cell tracks (Fig. 1g, and Supp. Fig. 11). To facilitate the visualization and analysis of these data; a *napari* plugin was developed that lets users select and follow groups of cells to determine their fate (Fig. 1h, Supp. Fig. 12 and Methods). We refer to this technique as *in silico* fate mapping. By conducting in silico fate mapping of a group of anterior-neural progenitors in the forebrain, we observed that their fate was limited to the forebrain region (Fig. 4a). This finding aligns with the RNA velocity dynamics of neural anterior state differentiation, as depicted by the purple star * in Fig. 3a. Furthermore, we investigated whether the well-known differentiation of mesodermal tissue visible in transcriptomic space corresponds to actual developmental lineages (Fig. 3a, green star *). We identified early presomitic mesoderm and observed clones that integrated into the somites, as expected (Fig. 4b, purple star *). All in all, our fate mapping analysis confirms cellular transitions identified in transcriptomic space.

### High-resolution time-resolved *in silico* fate mapping predicts time-dependent NMP fate restriction

We next directed our attention towards the late pluripotent progenitors NMPs (Fig. 3a, red star *). To determine the presumptive NMP domain, the spatiotemporal localization of NMPs was mapped by HCR (Hybridization Chain Reaction) based on their co-expression of *sox2* and *tbxta* (Supp. Fig. 13). The *in silico* fate mapping of the early NMP territory (<14 hpf) confirms that NMPs give rise to mesodermal and neural progeny (Fig. 5b, Supp. Fig. 14a and Supp. Video 7). The fraction of progeny was quantified, and 48% of cells went to the somites, 16% went to the neural tube, 28% went to the tail bud, and 8% went to the pre-somitic mesoderm (Fig. 4c and Supp. Fig. 14e). The ventrolateral domain posterior to the NMPs is the mesodermal progenitor territory (Supp. Fig. 14b, e and Supp. Video 8), the dorsomedial territory posterior to the NMP area is the neural progenitor zone (Supp. Fig. 14c, e and Supp. Video 9). These *in silico* fate maps support the established model of zebrafish posterior developmental specification domains^22, 42^ (Supp. Fig. 14 e). Interestingly, fate maps of NMPs changed at later time points (after 14 hpf, see Fig. 4d, Supp. Fig. 14 d and Supp. Video 10), where NMPs primarily contributed to mesodermal tissue. Most late NMP progeny integrated into the pre-somitic mesoderm and somitic mesodermal tissues (38% and 28%, respectively, see Supp. Fig. 14e), while 32% remained in the tail bud. In conclusion, the virtual fate mapping of axial progenitors indicates that NMPs are effectively pluripotent only during the beginning of axial elongation.

### *In vivo* fate-mapping via photo-conversion confirms time-dependent NMP fate restriction

To validate the results obtained with our *in silico* fate mapping experiments, we performed photoconversion followed by live light-sheet imaging of developing zebrafish embryos (Supp. Fig. 15, Supp. Fig. 16, and Methods). Using embryos ubiquitously expressing the photoconvertible protein Dendra2^43^, the presumptive territory of NMPs was photoconverted at different developmental stages based on previous observations (Supp. Fig. 13). During early segmentation (10 hpf – Fig. 5a and Supp. Video 11) progeny from the photoconverted NMP territory were distributed in mesodermal and neural tissues (Fig. 5b, c and Supp. Video 11). However, when the NMPs’ presumptive territory was photoconverted at 14 hpf (Fig. 5e and Supp. Video 12), the progeny only integrated into the mesoderm (Fig. 5f, g and Supp. Video 12). The distribution of the clones was computed by segmenting the 3D images and quantifying the NMP progeny in the manually annotated territory (mesodermal tissue and neural tissue, Fig. 5d, and h). Early photoconversion resulted in 46.2% of progeny in neural tissues and 53.8% of progeny in the mesodermal tissues (Fig. 5d). In contrast, photoconversion of NMPs at 14 hpf resulted only in 3.5% in the neural tube, and a large majority (96.5%) in the mesoderm (Fig. 5h).

### Manual track curation confirms time-dependent NMP fate restriction

To verify that NMPs are truly pluripotent cells versus just a combination of lineage-specific progenitors (i.e., ‘salt and pepper’ distribution), we tracked single photoconverted putative NMPs. Cells that integrated at later stages into the neural tube and mesoderm were automatically backtracked to search for a common ancestor originating in the presumptive NMP territory. Then the resulting tracks were manually curated – a tedious process because of tissue crowdedness and fast cell dynamics (Fig. 5i). We were able to reconstruct the track of an early NMP that eventually divided and which progeny integrated into both mesodermal and neural tissues (Fig. 5j and Supp. Video 12). However, when this search was repeated for late photoconversion experiments, no pluripotent cell originating from the presumptive NMP territory could be identified, confirming our *in silico* fate mapping results.

### Tissue morphodynamics constrains the fate of NMPs

We next tested whether spatial positions and distinct tissue deformation environments could affect NMPs’ fate specification. To this end, we applied the recently developed dynamic morphoskeletons (DMs) framework^44^. This framework aims to identify Lagrangian Coherent Structures (LCS), distinct structures in the embryonic cell flow that organize and compartmentalize cellular trajectories^44^. Of relevance for NMP specification are repelling LCSs (or simply repellers) that mark regions where cells distinctly separate over time. Repellers were identified by computing the Lagrangian stretching of cellular flows derived from the tracking data (Supp. Fig. 17a, Methods, and Supp. Information). The analysis revealed that from 10 to 12 hpf during the early stages, the NMPs (localized via HCR, Supp. Fig. 17b see Supp. Fig. 18) are located on a repeller (see Fig. 7b, c, and d). This repeller is characterized by neighboring cells rapidly separating from each other (Supp. Fig. 17a, see vectors Supp. Fig. 17c). The repelling structure was strongest between 12 and 14 hpf, a time during which the NMPs are effectively pluripotent (Supp. Fig. 17d - middle). This is consistent with the antagonist cell flows observed at the NMP territory (Supp. Fig. 17c, center). In contrast, no repelling structure was present during the following time window when the NMPs’ fate is restricted to the mesoderm (after 14 hpf, Supp. Fig. 17d - bottom). At that point, the cell flow is coherent and solely directed toward the mesoderm (Supp. Fig. 17c - bottom). Therefore, cells in that region are now effectively prevented from reaching the neural tube. The same was true for the subsequent time window (Supp. Fig. 19). These results suggest that tissue compartmentalization is correlated with, and can potentially explain, the observed fate restriction of the NMPs.

Using single-cell imaging-based lineage reconstruction, we observed a state transition of NMPs. Our findings revealed that during early gastrulation, NMPs exhibited characteristics of true bipotent progenitors, generating progeny that integrated both mesodermal and neural tissues. However, at mid-somitogenesis, the fate of NMPs is restricted to the mesodermal lineage. We went back to Zebrahub sequencing resources to characterize the transcriptomic landscape of NMPs.

### The transcriptomic landscape of NMPs’ *Sox2*/*tbxta* gradient predicts their future transcriptomic state

Next, we investigated the NMP’s transcriptomic state to find if it corresponds to the fate mapping temporality. We zoomed in on the NMPs and the adjacent clusters and re-projected the data on a new UMAP (Fig. 6a, Supp. Fig. 10d). In this projection; the NMPs stay positioned between the presomitic mesoderm (PSM) and neural tube (Fig. 6a and Supp. Fig. 20). The RNA velocity analysis on the new projection showed the same mesodermal and neural flows originating from the NMP population (Fig. 6b and Supp. Video 6). Flows, originating from the NMP cluster, were observed towards the notochord (mesodermal tissue) and floor plate (neural tissue - Fig. 6b) – two cell populations located along the midline of the vertebrate’s tail. Notably, we found a *sox2/tbxta* expression gradient for NMPs: cells expressing more *tbxta* were closer to mesodermal lineages, whereas cells expressing more *sox2* were closer to neural cells (Fig. 6c). This result suggests the importance of *tbxta* and *sox2* transcription factors in pushing the NMPs toward the mesodermal or neural fates, respectively.

### NMP population size decreases during early development

We quantified the relative proportion of sequenced NMP cells between 10 hpf and 24 hpf and observed a decrease over time with a more substantial decline between 12 hpf and 16 hpf (Fig. 6d). To corroborate this observation, Hybridization Chain Reaction (HCR, see Methods) was used to stain *sox2* and *tbxta* mRNA in fixed embryos. We then imaged the samples, segmented the cells, and counted those that co-expressed *sox2* and *tbxta* (Fig. 6e, f and Supp. Fig. 13a and Methods). This analysis was consistent with the scRNAseq-derived quantification, confirming a reduction of NMPs between 12 hpf and 16 hpf (Fig. 6g and Supp. Fig. 13b).

### Gene-expression dynamics suggest that early NMPs are pluripotent and late NMPs mainly contribute to the mesoderm

To better understand the temporal dynamics of NMPs during early development, RNA velocity data were analyzed at individual time points (10, 12, 14, 16, 19, and 24 hpf). At ten hpf (the end of gastrulation, beginning of axial extension – Fig. 6h), the NMP population appears closer to the posterior neural cluster with RNA velocities directed towards both mesodermal and neural clusters. A similar pattern was observed at 12 and 14 hpf: the two main RNA velocity flows originating from the NMPs are directed towards both PSM (mesoderm) and neural posterior clusters. However, at 16 hpf, a shift was observed: the NMP cluster attached only to the mesoderm with their RNA velocities extending exclusively toward the mesoderm. A similar configuration is kept at 19 hpf and 24 hpf, with the RNA velocity of NMPs directed toward the mesoderm. Interestingly, at almost every stage, the cell flow originating at the NMPs is also partly directed to the notochord (mesoderm) and a lesser extent, to the floor plate (neural). To quantify this observation and prevent ambiguities due to the UMAP projections, the distribution of RNA velocities for NMPs along the neural-mesodermal axis was computed at each developmental stage (Fig. 6i). This analysis confirmed that during early stages (10, 12, and 14 hpf), NMP velocities point towards both mesoderm and neural tissue. In contrast, they primarily point towards the mesoderm during later stages (16, 19, and 24 hpf).

### Pseudotime lineage reconstruction supports NMPs’ fate restriction

To further understand the developmental progression of NMPs along the neural-mesodermal axis, pseudotime developmental trajectories^45^ were reconstructed. These trajectories recapitulate the expression of genes that play crucial roles in neural and mesodermal differentiation^46^ (Fig. 6j). The critical cell types in these trajectories were also identified (Fig. 6k). Their distribution was visualized in developmental time (Fig. 6l). These findings were consistent with our previous observations, indicating that the majority of NMPs moving in the neural direction are early-stage cells, while the cells moving in the mesodermal direction are from all stages (Fig. 6k, l – red bar). Overall, both RNA velocity and pseudotime reconstructions suggest a fate transition for NMPs at 16 hpf: Early NMPs have the potential for both mesodermal and neural fates, whereas, after 16 hpf, they are restricted to the mesodermal lineage. Overall, the dynamic analysis of the transcriptomic space and the image-based lineage reconstruction strongly indicate a state transition of NMPs, shifting from bipotent progenitors to committed progenitors with a mesodermal fate.

### Dissecting the molecular dynamics of NMPs

Next, we investigated the expression dynamics of genes associated with molecular pathways that are important in axial development. Amongst the transcripts specifically enriched in NMPs (Supp. Table 2) were genes involved in the control of body elongation (*cdx1a*, *lin28a*, *cdx4, gdf11*) ^47–49^ as well as genes related to FGF, RA, and Wnt signaling pathways, important for mesodermal and neural induction^40^ (Supp. Fig. 21a). Interestingly, enriched genes showed a peak of expression at early stages of segmentation (Supp. Fig. 21a – 10, 12, 14 hpf). Remarkably, the well-known pluripotency factor *oct4*^50^ was among the enriched genes at 12 hpf (encoded by *pou5f1*, arrow in Supp. Fig. 21a). We confirmed the specific expression of *oct4* in NMPs by comparing it with other tissues and confirmed that after 10 hpf, *oct4* expression is maintained at a high level in NMPs compared to different cell types (Supp. Fig. 21b). We also plotted genes that are important for the cell cycle^51^ and observed an upregulation of these genes at 12 hpf, suggesting an increase in proliferation at this stage (Supp. Fig. 21c). We clustered the cells annotated as NMPs in a new UMAP and observed a small population of cells located between the early and the late cell clusters (Supp. Fig. 21d mixed population), with a high expression of *tbxta*/*sox2* and enriched for genes specific to the notochord lineages (*noto*, *ntd5*, *shha*). This observation echoes studies describing a population of progenitors in a vertebrate’s posterior territory contributing to the notochord^52–54^. As expected, we observed a striking co-linear activation of the hox genes indicating a temporal maturation of the NMPs as the body extends (Supp. Fig. 21e). Taken together, our results suggest that NMPs are enriched in pro-mesodermal, pro-neural genes pathways and a pluripotency factor with the highest expression during early segmentation.

### Interactive data sharing at zebrahub.org

To ensure widespread access to the Zebrahub datasets, protocols, software, and instruments, we have developed an interactive website (Fig. 7a). Users can access the scRNAseq datasets and light-sheet microscope time-lapse within their web browsers (Fig. 7b). The scRNAseq datasets can be visualized in an interactive tool, allowing users to explore metadata, annotations, search for specific genes, and create gene sets (Fig. 7c). The processed light-sheet imaging time-lapse videos are easily accessible, enabling users to visualize volumetric datasets over time through various ortho-views (Fig. 7d). The *in silico* fate mapping software is downloadable alongside different datasets, with clear instructions on how to run and utilize them. This user-friendly and comprehensive resource bridges the gap between technical knowledge and domain expertise, facilitating the exploration of complex high-dimensional datasets and facilitating the discovery process.

## DISCUSSION

We introduced Zebrahub, a foundational multimodal resource for the comprehensive analysis of zebrafish development. It consists of single-embryo single-cell temporally resolved transcriptomes and light-sheet imaging datasets. All datasets, designs, and software are accessible for download. Sequencing, and imaging datasets can be explored interactively, and image-based lineages can be visualized and queried using our *napari*^55^ plugins. Zebrahub was created thanks to a wide range of multidisciplinary advancements, such as a protocol for high-efficiency single-embryo cell dissociation, a framework for studying the transcriptomic variability of individual embryos, an open-source design for adaptive multi-view light-sheet imaging and photo-manipulation, as well as computational tools for *in silico* fate mapping from light-sheet imaging data.

Furthermore, in contrast to previous scRNAseq studies^11–16, 19^, our resource encompasses a broader developmental period, attains non-pooled resolution at the single-embryo level, and our gene coverage per embryo is sufficient to measure transcriptome variability between individuals. In addition, we used state-of-the-art technology, including a high-resolution (1.0 NA) and large-volume (> 1mm^3^) light-sheet microscope^33^, automated whole embryo cell tracking, and a theoretical framework^44^, to investigate the lineages and cellular movements during late stages of vertebrate embryonic development. We could achieve this in highly dynamic and densely populated tissues like those in the developing tail. Interestingly, we observed discrepancies between the known developmental trajectories of specific cell types and their corresponding predicted cellular state progression in transcriptomic space (see Fig. 3d and results). This observation may be attributed to the inherent limitations of RNA velocity in accurately reconstructing cellular dynamics based on scRNAseq data^56^, or it could be affected by the resolution of the clusters used for annotation. Higher resolution annotations over more specific cell types could help clarify this. However, cell identity is still highly malleable at early stages because the spatial context influences cellular signaling and, consequently, cell fate. Nevertheless, these observations

At the tissue level, we observed a repeller structure indicating rapidly diverging cell flows only in the early NMPs (before 15 hpf), supporting that NMP progeny integrates two different tissues (Supp. Fig. 17). In contrast, when the axial progenitor territory is more stable (no repeller), and cell flow is coordinated, the NMPs remain pluripotent competent, but give only rise to mesoderm. Interestingly, the RNA velocity study also suggests that the NMPs might be at the origin of other axial tissues, namely notochord and floor plate. However, in the image-based fate mapping analysis, there is no clear evidence of this relationship (Fig. 4 and Fig. 5). Hence, the crucial role of multimodal fate mapping atlases like Zebrahub for understanding the complex dynamics occurring during embryogenesis and reconciling spatial and transcriptomic insights on cell fate and lineage specification.

Zebrahub’s collection of readily accessible datasets and tools represents a foundational resource to the scientific community. By combining imaging and sequencing datasets, we joined the strengths of both modalities. Besides NMPs, this resource will shed light on other important developmental processes. Zebrahub is poised to expand with more developmental stages, novel multiomic datasets, and ultimately a consensus digital lineage reconstruction of multiple embryos. Integrating existing datasets, including from other species, will enable the creation of a comprehensive atlas of vertebrate embryogenesis, ushering in a new era for developmental and evolutionary biology.

## Supporting information

Supplementary Video 3. Light-sheet time-lapse of Tg(h2afva:h2afva-mcherry; mezzo:eGFP) embryo starting at 50% epiboly.

Supplementary Video 4. Image processing and analysis steps for nuclei segmentation and tracking.

Supplementary Video 5. Dynamic visualization of the RNA velocity flows overlayed on full UMAP color-coded by annotations.

Supplementary Video 6. RNA velocity-derived flows around the NMP cluster

Supplementary Video 7. In silico fate mapping of the early NMP presumptive territory.

Supplementary Video 8. In silico fate mapping of the mesodermal axial progenitors.

Supplementary Video 9. In silico fate mapping of the neural axial progenitors.

Supplementary Video 10. In silico fate mapping of the late NMP presumptive territory.

Supplementary Video 11. In vivo Photoconversion of the early NMP presumptive territory using the tg(bactin2:memb-Cerulean-2A-H2B- Dendra2) transgenic

Supplementary Video 12. In vivo Photoconversion of the late NMP presumptive territory using the tg(bactin2:memb-Cerulean-2A-H2B- Dendra2) transgenic l

Supplementary Video 13. Tracking of an NMP with daughter cells integrating both mesodermal and neural tissues.

Supplementary Table 1. Annotation ontology tracker.

Supplementary Table 2. Differential gene expression for annotated cells at 10, 12, 14, 16, 19, and 24 hpf.

## ACKNOWLEDGEMENTS

We thank the whole Data Science and IT teams of the Chan Zuckerberg Biohub for helpful discussion and support. We are grateful for the support, feedback, and proofreading from S. Schmid, J. DeRisi, and S. Quake. We thank P. Keller for the critical reading of the manuscript and help with our variant of the SiMView microscope design. We thank C. Guillot, L. Mahadevan, and S. Darmanis for initial discussions. We thank J. Weissman for supporting and encouraging the pursuit of zebrafish as the lab’s model organism. We thank A. York and Calico Life Science LLC for developing and loaning the AMS-AGY V2.0 objective. We also would like to thank J. Maitin-Shepard and V. Jain for their generous help with Neuroglancer. Funding for this work was provided by Chan Zuckerberg Biohub San Francisco (CZB SF). We thank the CZB SF donors, Priscilla Chan and Mark Zuckerberg, for their generous support.

## AUTHOR CONTRIBUTIONS

M.L. and L.A.R. conceived the project and designed the experiments. L.A.R. supervised the project. M.L. led the experimental and computational work. S.V., M.B., and M.L. developed the cell dissociation protocol and performed the library prep. S.V. and M.L. performed the HCR experiments. A.De., S.P. and H.M. performed the sequencing. A.G., S.A., A.M., and A.O.P. performed computational analysis for scRNAseq data. A.G. and S.A. designed the single fish computational pipeline. A.G. performed the cell cluster transition graph. M.L., T.L., R.B., A.J., K.B., S.V. and X.Z. annotated the cells. B.Y., Y.L., and H.K. developed and built the microscopes. M.L. performed light-sheet imaging experiments. S.V. performed confocal imaging experiments with help from M.L.; J.B., A.C.S., and L.A.R. wrote the light-sheet image processing code. R.G.S. designed and built the imaging chamber. J.B. developed the *in silico* fate mapping software. R.A., H.K., and L.A.R. wrote the code for microscope control. M.S. and S.S. wrote the code for the dynamic morphoskeletons framework and performed the analysis with J.B., M.L., and L.A.R. J.B. analyzed the imaging data. M.L., A.G., S.A., A.O.S., and L.A.R. analyzed the sequencing data. K.A., S.D., A.C.S., and L.A.R. developed the data portal. O.P. and G.H. provided feedback on the project conception. A.O.S., N.N., M.L., and L.A.R. supervised method development. A.Di., M.L., J.B., and L.A.R. generated the 3D visualizations for imaging and scRNAseq data. M.L. drafted the first version of the manuscript with input from A.G. and S.A., and M.L. and L.A.R. wrote the final version with feedback from all co-authors.

## CONTACT FOR REAGENT AND RESOURCE SHARING

Further information and requests for resources and reagents should be directed to and will be fulfilled by Loic A. Royer.

## DATA AVAILABILITY

Raw sequencing data are available at NCBI’s SRA BioProject PRJNA940501. Raw imaging data are available at zebrahub.org.

## CODE AVAILABILITY

All relevant code repositories are listed on zebrahub.org.

## INSTRUMENT AVAILABILITY

Blueprints, parts lists, control codes, and instructions can be found at repositories listed on zebrahub.org.

## METHODS

### Animals

The care and experimental procedures for zebrafish complied with the protocols approved by the institutional animal care and use committee at the University of California San Francisco (UCSF). The fish were bred and maintained at 28.5°C and were fed thrice daily using an automatic feeder^60^. Embryos were raised at 28.5°C and staged based on hours post fertilization (hpf), with the number of somites used as temporal landmarks^61^ during somitogenesis. We employed wild-type EKW embryos for the scRNAseq study. For the light-sheet experiments, we used either the tg(*h2afva:h2afva*-mcherry)^34^ line or the same line crossed with tg(*mezzo*:eGFP)^32^ (a gift from J. Huisken, University of Gottingen). The tg(*bactin2*:memb-Cerulean-2A-*H2B*-Dendra2) line^43^ was used for the photo-conversion experiments (obtained from the European Zebrafish Resource Center).

### Hybridization Chain Reaction (HCR) and imaging

We investigated the spatiotemporal localization of *sox2* and *tbxta* mRNA transcripts across six developmental stages (10, 12, 14, 16, 19 hpf, and 1 dpf). A probe set designed for the target mRNAs was obtained from Molecular Instruments. To prepare the embryos for *in situ* hybridization, we dechorionated them using a 1mg/ml pronase solution for no longer than 90 seconds, then washed them thoroughly with egg water. Next, the embryos were fixed with 4% paraformaldehyde solution overnight at 4°C. We performed RNA *in situ* hybridization according to the Molecular Instruments RNA FISH protocol for zebrafish embryos (MI-Protocol-RNAFISH-Zebrafish-Rev9, adapted from Choi et al.^62^) using the probe set. Following amplification, the embryos were incubated with DAPI (1:3000 dilution) for 20 minutes during the first wash step and gradually transferred to an 80% glycerol solution for storage. Finally, the embryos were carefully mounted whole in 80% glycerol on a glass slide and imaged using a confocal microscope (Andor Dragonfly).

### HCR image processing and quantification

We utilized the HCR images to quantify the proportion of presumptive NMPs based on the co-expression of *sox2* and *tbxta* mRNA transcripts. The images were first manually cropped to isolate the region of interest. Next, we denoised the images using <u>Aydin</u>’s auto-tuned Butterworth filter. Aydin^63^ is a powerful and user-friendly image-denoising tool that can significantly improve the image quality of microscopy datasets (http://aydin.app). Images were then downscaled along the z-axis by a factor of two to obtain isotropic images. The cells were then segmented using the DAPI channel in two steps. Firstly, we detected cells against the background and split the detected regions into individual cell instances. For the first step, we filtered the DAPI channel using a morphological opening operator^64^ and then removed the gaps between nuclei using a morphological area-closing operator. We then thresholded the images using the Otsu method to identify cells against the background. In the second step, individual cells were split using the previously opened images, which were inverted, normalized, and provided as input to the watershed algorithm^65^.

### NMP cell counts from HCR images

Before the gene expression measurement of individual cells was extracted, we homogenized the illumination of the z-slices. For every channel, each z-slice of each image was corrected using the DAPI channel as a reference, and the background fluorescence was removed using an area-opening morphological operator per z-slice. We then extracted the total intensities of the *sox2* and *tbxta* probes for each cell and classified these as presumptive NMP by manually adjusting the threshold. The threshold was chosen empirically to include all visible mRNA transcripts while avoiding background and fluorescence artifacts.

### Single-embryo cell dissociation

For scRNAseq experiments, a single embryo was used per experiment, with four replicate experiments (siblings) per developmental stage. The cell dissociation protocol is summarized in Supp. Figure 1. First, embryos were dechorionated manually with forceps. Next, a single embryo of the required stage was dissociated in either 50 μl of FACSmax solution (Gentalis - T200100) for embryos between 10 hpf to 1 dpf, or 50 μl of 0.25% Trypsin-collagenase (100 mg/ml) solution for embryos between 2 dpf to 10 dpf. The dissociation was performed by gently pipetting and flicking the Eppendorf to prevent bubble formation. The resulting solution was centrifuged at speeds varying gradually based on the stage of interest, with lower speeds of 300 rcf for 2 mins being used for earlier stages and higher speeds up to 750 rcf for 7 mins for later stages, all while maintaining a temperature of 4°C. The dissociation media was carefully removed, and the pellet was washed with 1X PBS + 1% BSA solution. For embryos between 2 to 10 dpf, the solution was passed through a 20 μm mesh to remove undissociated debris, and then the sample was centrifuged again under the same conditions at 4°C. The supernatant was discarded, and the pellet was dissolved in 25 μl of 1X PBS+ 0.1% BSA + 18% optiprep. The cell concentration was measured using a cellometer auto T4 cell counter (Nexcelom Biosciences). The cell count was determined using a dilution (1:4) of the cell suspension and trypan blue solution.

### Library preparation and sequencing

The single-cell suspensions were processed into droplet emulsions using the Chromium Single Cell Controller (10x Genomics), and scRNA-seq libraries were generated with the Chromium Next GEM Single Cell 3’ Reagent Kit v3.1 (10 hpf to 1 dpf) and Single Cell 3’ Reagent Kit v3.1 HT (2dpf to 10 dpf). Before loading single-cell GEMs, the single-cell suspensions were assessed using the Cellometer Auto T4 Cell Counter to determine the optimal cell count, which ranged from 700 to 1200 cells/ul for loading onto the Chromium Controller. Single cells were loaded into each channel with the appropriate amount of diluted RT master mix based on the measured cell concentration and targeted cell recovery per sample. Fragmentation, ligation, and PCR amplification were performed using the BioRad C1000 Touch Thermal Cycler with thermal profiles specified by the 10X Genomics User Guide CG000204. Amplified cDNA and final libraries were assessed using the Agilent 4200 TapeStation/High Sensitivity D5000 Screentape and qPCR with the Kapa Library Quantification kit for Illumina. Each library was normalized to 4 nM and pooled for each NovaSeq sequencing run. The pools were sequenced with Novaseq run kits, using either single or dual index plates (Illumina 20028313), with a 1% PhiX control library spiked in. Sequencing was performed on the NovaSeq 6000 Sequencing System (Illumina).

### Read alignment and single-ell filtering

After sequencing, the obtained sequences were de-multiplexed using <u>bcl2fastq</u> (v2.20.0.422). To align the reads to the reference genome, a reference genome was first created using <u>CellRanger</u> v.4.0.0. Subsequently, the CellRanger v.5.0.1 software was used for read alignment to the reference zebrafish genome.

### Filtering and clustering of single-cell RNA sequencing data

We performed quality control to filter out low-quality cells by removing cells with counts below 10,000 or with a fraction of mitochondrial genes greater than 5%. We then identified 2000 highly variable genes for each time point using Scanpy (v1.9, see <u>scanpy.readthedocs.io/</u>) and conducted Principal Component Analysis (PCA). We selected the first 100 principal components to cluster the single cells. To visualize the cells in a two-dimensional space, we applied the UMAP algorithm^28^ (see <u>umap-learn.readthedocs.io</u>) to explore lower-dimensional representations of our gene expression data and facilitate the annotation of cell types. The UMAP projection showed that data from different embryos did not exhibit significant batch effects, as shown in Fig. 3.

### Integration of multiple time points

To create a comprehensive UMAP embedding that included all time points, we employed the integration pipeline from Seurat^66^ (v3, see satijalab.org/seurat/), a software tool used for single-cell genomics. Typically, data integration pipelines are utilized to rectify batch effects, which can lead to an over-correction of actual biological differences. However, we anticipate that some differences between time points would correspond to genuine biological changes during development. To prevent overcorrection, we utilized the integration pipeline with reciprocal PCA. This procedure generated a PCA projection of all cells from all time points. Finally, we used the first 30 principal components of the integrated PCA space to produce 2-dimensional and 3-dimensional projections using the UMAP algorithm (as depicted in Fig. 1e, Supp. Video 2, and 3).

### Clusters annotation. We chose a higher clustering resolution to ensure a comprehensive representation of cellular diversity

than is typically required. We identified cell types based on the expression of cluster-specific enriched markers obtained from literature search and ZFIN database queries (see zfin.org). For each developmental stage, individual UMAPs were annotated using a primary zebrafish expert. Then they verified the annotations independently with a second expert—a third expert used zebrafish ontology terms from the database to homogenize cell-type naming across all time points. We reannotated cell clusters following the same procedure for the integrated UMAP centered around the NMPs population. To aid interpretation, we also used coarser annotations of broader cell types, which are summarized in Supp. Table 1.

### Gene expression variance across sibling embryos

We devised a framework to quantify gene expression differences across individual embryos. Our framework relies on estimating probability distributions of gene expression across single cells for each embryo and applying an information-theoretic approach to quantify the differences between those distributions. First, for each gene, we took the log normalized counts and grouped the counts by embryo. We filtered out genes with low counts; specifically, we discarded genes for which 80% of the cells had zero counts. We also removed genes with a mean expression less than 0.1 to prevent skewed distributions towards zero values. We also excluded genes with less than ten non-zero counts for each embryo. After quality control, we fitted a probability density function for the expression of each gene (n = 4 embryos per time point) using a Gaussian kernel (function scipy.stats.gaussian_kde v. 1.9.1, with the default ‘scott’ method to calculate the estimator bandwidth, see Fig. 2c, d, and Supp. Fig. 7). Finally, we used the Kullback-Leibler divergence to estimate, for each gene and per stage, all pairwise divergences between embryo gene count distributions (scipy.stats.entropy v. 1.9.1, and see Fig. 2c, d heatmaps, and Supp. Fig. 7). The KL divergence is a statistical distance that measures the difference between two probability distributions over the same range of values. We sampled 1000 values over the same range (min(x,y) - 1, max(x,y) + 1) for each pair of distributions P(x), P(y). Since the KL divergence is not symmetric, we computed the pairwise distances in both directions (KL(Px,Py) and KL(Py,Px), for x**≠**y, n = 4^2^-4 values). We then summarized all 12 pairwise KL scores with the median to get the mpKLD. Repeating this process for all genes, we obtained a distribution of mpKLDs for the whole transcriptome. We performed this analysis on different cell populations, i.e., for cell types and the entire embryo. The study was conducted independently at each time point. We only analyzed cell types with at least 80 cells per embryo to ensure our distribution estimates were robust.

### A null model for median pairwise KL divergences (mpKLD) through embryo ID randomization

To establish a negative control and obtain a baseline for differences in gene expression between embryos, we performed a bootstrap-like method by randomly reassigning the embryo-specific ID for each cell. We thus obtained expected mpKLD values as if the differences between embryos were due to chance alone. We repeated this process 20 times, which produced the expected mpKLD values for each gene based on the randomized embryo IDs (see Supp. Fig. 8a, red and blue histograms). To evaluate the significance of the observed differences between embryos, we calculated a z-score for each gene by comparing the observed distribution of mpKLD values to the scrambled distribution. We observed a robust linear relationship between the mpKLD scores and the z-scores (see Supp. Fig. 8a, scatterplot). This null model analysis allowed us to assess the significance of observed differences in gene expression between individual embryos.

### RNA velocity

RNA velocity is a method that estimates the rate and direction of changes in gene expression for individual cells. We used scVelo (v0.2.5, <u>scvelo.readthedocs.io</u>) to perform RNA velocity analysis. We started with the CellRanger alignment and identified spliced and unspliced variants for all genes using Velocyto^39^ (v0.17, see velocyto.org). Next, using scVelo’s dynamic modeling approach^67^, we computed the RNA velocity for each gene and cell. This lets us estimate the rate and direction of changes in gene expression for each cell in our dataset. Finally, we projected the average velocity vectors as a stream plot onto the integrated UMAP to visualize the changes in gene expression over time. We repeated this process for each time point to investigate how RNA velocity vectors change over time. The resulting analysis is presented in Figure 3b.

### RNA velocity graph

The RNA velocity analysis allowed us to obtain a probability transition matrix between single-cell states, which describes the likelihood of a cell transitioning from one state to another based on the dynamics of gene expression derived from RNA velocity. To construct this matrix, we used the kernel function provided by CellRank^41^ (see <u>cellrank.readthedocs.io</u>), which considers both the velocity vectors and the similarity between single cells. The resulting matrix and its associated graph can be used to define a Markov process^41, 67^, which lets us estimate the transition rates between individual cell states and identify potential developmental trajectories. In other words, this approach models the temporal dynamics of gene expression in individual cells and predicts their transcriptional changes as they differentiate and as the tissue develops over time. The resulting graph can be visualized to provide a comprehensive overview of the transcriptional trajectories of cells during development.

### Cell cluster transition graph

We assumed that developmental dynamics follow a Markov process such that the weighted graph defines the stochastic probability matrix^41^. To calculate the transitions between cell clusters, we considered independent realizations of the Markov process where each cell cluster (cell state or type) is fixed as the initial condition. Progenitor clusters, for example, will transition into their corresponding intermediate states, whereas terminal populations will not. Therefore, we defined the initial states of the Markov process such that all cells in each cluster start with probability 1/n, where n is the number of cells in the cluster. By applying iterative multiplication of the stochastic matrix, we obtained the distributions of cell fates at different steps. At each Markov step, we sum up the probabilities of single cells within each cluster to get average cell state probabilities (Fig. 3b) and repeat this process for all 39 cell clusters. To generate a coarse-grained velocity graph, we defined edges as the immediate transitions between cell clusters (Fig. 3b – velocity figure cartoon with dynamics) according to their average transition probabilities within the first 10-time steps, when most cell states have left their initial state (except when they are terminal).

### RNA velocity quantification along the mesodermal-neural axis

We used the velocity graph as previously described. For each NMP cell, we computed the graph’s weighted sum of outcoming velocity magnitudes. Finally, to distinguish between the neuronal and mesodermal lineages, we defined velocity magnitudes toward neurons as “negative” and those towards mesoderm as “positive” and plotted the distribution of velocity magnitudes per time point for all NMPs (Fig 4i).

### Pseudo-time trajectories

Based on the integrated embedding (Fig. 4a), we found potential developmental trajectories for NMPs using Slingshot^45^ (see github.com/kstreet13/slingshot). Briefly, Slingshot fits a principal curve in a high-dimensional space such that single cells map to a one-dimensional projection called pseudo-time. As input for Slingshot, we used a 3-dimensional UMAP projection of the data and defined the root as the cluster with the highest number of NMPs. In agreement with our results from RNA velocity, Slingshot identified two main trajectories, one towards mesodermal cell types and another towards neuronal cell types. We verified that these two trajectories comprised the expected cell populations with the corresponding differentiation markers (Fig. 3j, k). As a control, we checked the correspondence of pseudo-time to real-time and verified that the most differentiated cells present at the end of the trajectories came from the latest time-point (24 hpf, see Fig. 3l). Finally, to find differential gene expression across pseudo time, we used tradeSeq^68^ (see github.com/statOmics/tradeSeq) and tested for differences between the neuronal and mesodermal lineages. We also found genes specifically associated with each lineage (Fig. 3j).

### Aligned UMAPs of individual time points

The stochastic nature of the UMAP algorithm leads to variations in the orientation, scale, and layout of UMAP projections. This impedes the comparison of datasets, such as developmental stages (e.g., Fig. 3h), as it is challenging to distinguish between (i) genuine differences in the data and (ii) differences caused by algorithmic stochasticity. **We aligned projections by utilizing pairwise anchors between different datasets to address this issue**. For each point (cell), the nearest neighbors in the previous and next developmental stages were used as anchors. To prevent over-alignment artifacts, we relaxed the strength of the alignment regularization parameter to 0.001. The outcome is a series of UMAP projections, each corresponding to a time point, that exhibit conserved relative orientation and proportions with one another (Fig. 3h). We confirmed that the aligned UMAPs maintained the biological relationships of known lineages as indicated by the expression of known markers and the distribution of cell type annotations (Fig 3h).

### Adaptive multi-view multi-color light-sheet imaging and processing

The open-source repository at github.com/royerlab/opensimview provides all information for building a two-detection and two illumination simultaneous adaptive multi-view multi-color microscope^30, 31^. We provide all blueprints, a parts list, the control software, and image processing tools. Once acquired, the dataset consists of multiple fused, deconvolved, and stabilized views. This processing was done with DEXP^69^ (see github.com/royerlab/dexp). DEXP is an image processing and visualization library based on <u>napari</u>, <u>CuPy</u>, <u>Zarr</u>, and <u>DASK</u> designed for managing and processing light-sheet microscopy datasets. It consists of light-sheet imaging specialized image processing functions (equalization, denoising, dehazing, registration, fusion, stabilization, deskewing, deconvolution), visualization functions (napari-based viewing, 2D/3D rendering, video compositing and resizing, mp4 generation), as well as dataset management functions (copy, crop, concatenation, tiff conversion). Almost all functions are GPU accelerated via <u>CuPy</u> but also have a <u>numpy</u>-based fallback option for testing on small datasets. In addition to a functional API, DEXP offers a command line interface that makes it very easy for non-coders to pipeline large processing jobs from a large multi-terabyte raw dataset to fully processed and rendered video in MP4 format.

### Sample preparation for multi-view light-sheet Imaging

Embryos were first gently dechorionated with sharp forceps under a binocular dissecting microscope. Embryos were gently transferred in FEP tubes (Zeus Inc. custom order) embedded in 0.1% low-gelling-temperature agarose (Sigma, A0701). The FEP tubes were placed in a custom-made sample capillary holder previously filled with 0.7% low-gelling temperature agarose to support the embryo and then placed in the chamber using a holder insertion tool and section pipette tip (sealed with low toxicity silicone – Kwik-Sil^tm^ from World Precision Instrument). A detailed procedure for the mounting is shown in Supp. Fig. 6a. CAD drawings of the custom-made FEP holder and the mounting chamber are available in Supp. Fig. 5. For recording after 15 hpf, fresh and filtered embryo water was circulated in the chamber with 0.016% tricaine.

### Photo-conversion experiments

For the photo-conversion, we used a python-based custom-made software (see photo-conversion arm in github.com/royerlab/opensimview) that allows the users to select a region of interest and photoconvert this area by controlling a two-axis galvo mirror actuator. We selected the embryo area to photoconvert based on our HCR mapping of the presumptive NMP territory (Supp. Fig. 13). Photoconversion was achieved using a 405 nm laser. Time and laser power were manually accessed based on the sample fluorophore state transition evaluation.

### Photo-conversion image analysis and quantification

**Estimating** the proportion of neural and mesodermal nuclei on the photo-manipulated light-sheet data requires three steps: segmentation of the nuclei; selection of which nuclei were photo-manipulated; nuclei label assignment to neural or mesodermal tissue. Before segmenting the nuclei, for each time-lapse, an individual frame (i.e., a 3D image) was selected at the time point of interest. The photo-manipulated channel fluorescence can be weak in some nuclei; thus, we used both channels (red and green) for segmentation. We removed the background using morphological filtering and thresholding. The images were normalized, so their intensities were between 0 and 1. These normalized images were forwarded into a custom-trained 3D U-Net^70^, where their nuclei and boundaries were detected. The nuclei detections were binarized with a threshold of 0.5, and the green and red channel detections were combined using a logical OR operator; the boundaries detection were combined using addition, resulting in a single binary nuclei detection and boundary per image. Finally, the nuclei were segmented using the hierarchical watershed^71^ and selecting the segments by picking a single level of the hierarchy (i.e., hierarchy horizontal cut). We extracted the original images’ total intensity of red and green fluorescence for each segment. The photo-manipulated nuclei were selected by manual thresholding of their total red fluorescence intensity – a single threshold per image. The regions of interest of each image were manually annotated into mesodermal or neural regions using napari^55^; these annotations were propagated into overlapping nuclei, assigning their respective labels to the nuclei’s segments.

### High-resolution & large volume single-objective light-sheet imaging

We used our DaXi imaging platform^33^ to perform high-resolution tail development imaging (1.0 NA). To improve coverage, we use two orthogonal, oblique light sheets for each time point. Images were processed using <u>DEXP</u>^69^ to (i) equalize brightness between views, (ii) remove background, (iii) correct for distortions between views, and (iv) temporally stabilize the timelapse to remove artifacts caused by sample movement or vibrations. Instructions to build and operate this instrument can be found are: github.com/royerlab/daxi

### Sample preparation for single objective light-sheet microscopy (DaXi)

Embryos are first dechorionated in a glass-bottom cell culture dish (35mm Cell Culture Dish with Glass Bottom 20mm – Stellar Scientific) and then gently transferred into a 0.7% solution of low-gelling-temperature agarose (Sigma, A0701). Embryos are then positioned at the correct position and angle for imaging using a custom-made capillary. When solid, the agarose surrounding the tail was cut off and removed using a dissection knife and forceps to permit full development and tail elongation (Supp. Fig. 6b). The time lapses were done with a gentle flow of fresh and filtered embryo-medium solution with 0.016% tricaine to prevent embryo movement.

### Whole embryo cell segmentation and tracking

As with the photo-converted images from the multi-view microscope, the images from the single-objective microscope were fused and stabilized using <u>DEXP</u>^69^. Once processed, the time-lapse was cropped to select the region of interest encompassing the embryo’s tail. Once again, the cropped video was stabilized to reduce further the local movement between frames of the highly dynamic process of tail development. Subsequent analyses were conducted on this cropped and stabilized time-lapse. To obtain the tracks, we used a hybrid approach combining multiple segmentation hypothesis tracking and the latest fluorescence imaging nuclei segmentation development using deep learning. We detect the nuclei and their boundaries using a 3D U-Net^70^ and the watershed algorithm^71^ to obtain the candidate segments. Instead of performing a horizontal cut to select segments, we use segment hierarchies to generate multiple segmentation hypotheses and let the tracking algorithm pick the most likely segments.

### *In silico* fate mapping

Despite recent advancements^72, 73^ cell tracking still needs to be solved and still requires extensive manual curation. Unfortunately, manual curation of tens or hundreds of thousands of cell trajectories is infeasible. We estimate the local cell flow between frames to circumvent this issue and use this as a robust alternative for analyzing the cells’ movements. For the local movement estimation, we use radial neighborhood regression^74^. For each time point, we fit a regression model that predicts the positions of cells in subsequent frames. The training data for these models are pairs of adjacent same-track coordinates at the time point of interest. Track endpoints (splits) are connected to the nearest starting point on the next frame. With these regression models, one for each time point, we can evaluate the movement of any cell at any location if they fall within the radius of an existing track. The tool is available at github.com/royerlab/in-silico-fate-mapping, and lets users predict the cells’ fate by estimating their movement over time given a user-selected starting point. This is analogous to a photo-manipulation experiment but *in silico*.

### Dynamic Morphoskeleton

To investigate cellular flows during morphogenesis in the zebrafish tailbud, we used the Dynamic Morphoskeletons (DM) method^44^, based on the Lagrangian Coherent Structures (LCS) concept. The DM is a self-consistent and objective method that identifies repellers, attractors, and their domains along cell trajectories by computing the Lagrangian stretching. The Lagrangian stretching is computed from cell trajectories using a recently developed technique that identifies LCSs from sparse and noisy trajectories^75^. For all results presented in this study, we set the radius of neighboring trajectories to a fixed value and used a time interval of interest. The Lagrangian stretching facilitates the identification of repelling spatiotemporal regions, which are marked by high values of Lagrangian stretching. It enables the characterization of differential mechanical environments for cell fate specification. The DM method is ideal for quantifying morphogenesis as it reveals the intrinsic geometric feature of spatiotemporal trajectories and is robust to noise. For more details explanation, see the Supp. Theory Note.

### 3D visualization & rendering The datasets’ visualization, interaction, and figure-making

were done using napari^55^ (see napari.org), particularly the napari-animation plugin (github.com/napari/napari-animation). We also used rendering routines in <u>DEXP</u>, some of which use the Spimagine (see github.com/maweigert/spimagine) volume rendering code. Moreover, the light-sheet datasets were stored using zarr^76^ to enable out-of-memory loading. 3D RNA velocity visualizations were rendered using the Houdini software.

## SUPPLEMENTARY VIDEOS CAPTIONS

[Supplementary Video 1. Zebrahub’s full dataset scRNAseq UMAP color-coded by annotations](https://vimeo.com/803717897/4a8cd4e7bb).

[Supplementary Video 2. Zebrahub’s full dataset scRNAseq UMAP color-coded by time points](https://vimeo.com/803717897/4a8cd4e7bb).

[Supplementary Video 3. Light-sheet time-lapse of Tg(*h2afva:h2afva*-mcherry*; mezzo*:eGFP) embryo starting at 50% epiboly](https://vimeo.com/803717897/4a8cd4e7bb).

[Supplementary Video 4. Image processing and analysis steps for nuclei segmentation and tracking](https://vimeo.com/803717897/4a8cd4e7bb).

[Supplementary Video 6. RNA velocity-derived flows around the NMP cluster](https://vimeo.com/803717897/4a8cd4e7bb)

[Supplementary Video 7. *In silico* fate mapping of the early NMP presumptive territory](https://vimeo.com/803717897/4a8cd4e7bb).

[Supplementary Video 8. *In silico* fate mapping of the mesodermal axial progenitors](https://vimeo.com/803717897/4a8cd4e7bb).

[Supplementary Video 9. *In silico* fate mapping of the neural axial progenitors](https://vimeo.com/803717897/4a8cd4e7bb).

[Supplementary Video 10. *In silico* fate mapping of the late NMP presumptive territory](https://vimeo.com/803717897/4a8cd4e7bb).

[Supplementary Video 11. *In vivo* Photoconversion of the early NMP presumptive territory using the tg(*bactin2*:memb-Cerulean-2A-*H2B*-Dendra2) transgenic line](https://vimeo.com/803717897/4a8cd4e7bb).

[Supplementary Video 12. *In vivo* Photoconversion of the late NMP presumptive territory using the tg(*bactin2*:memb-Cerulean-2A-*H2B*-Dendra2) transgenic line](https://vimeo.com/803717897/4a8cd4e7bb).

## SUPPLEMENTARY TABLE CAPTIONS

**Supplementary Table 1.** Annotation ontology tracker.

**Supplementary Table 2.** Differential gene expression for annotated cells at 10, 12, 14, 16, 19, and 24 hpf.

## SUPPLEMENTARY FIGURES

**Supplementary Figure 1.**
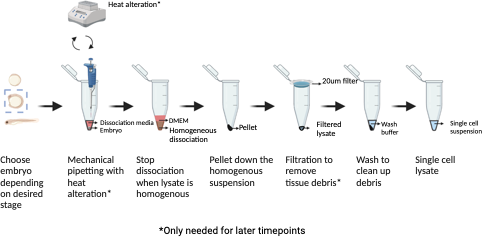
Schematic representation of the single embryo dissociation protocol for zebrafish embryos.

**Supplementary Figure 2.**
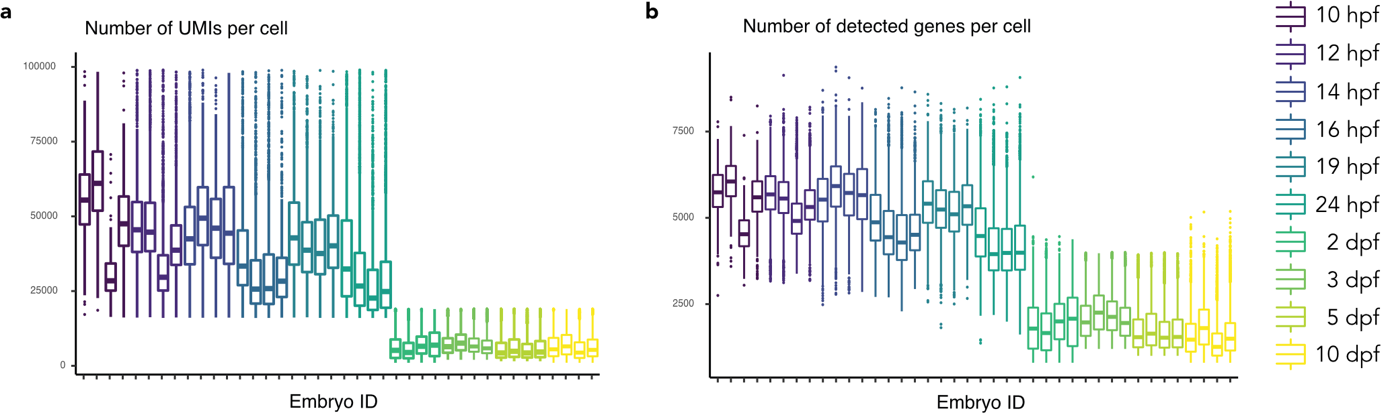
Distribution of UMIs and detected genes per embryo. The number of detected UMIs per cell for each embryo. Individual embryos are shown in the x-axis sorted by developmental time point (n=40 embryos, 10 developmental time points). The distribution of UMIs per cell is shown as a boxplot. An equivalent plot for the number of detected genes per cell.

**Supplementary Figure 3.**
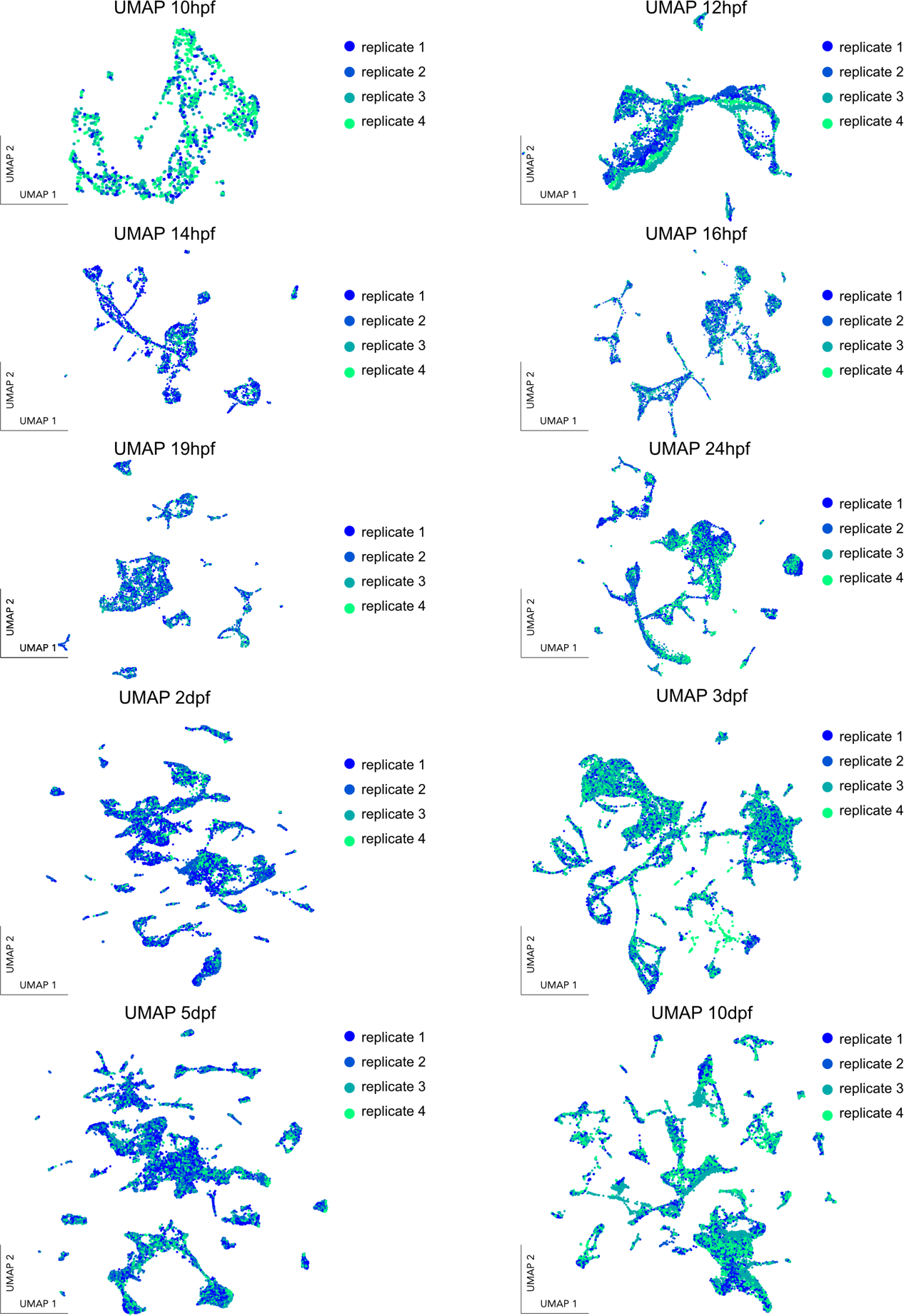
UMAPs for each timepoint colored coded by the individual replicate sequenced embryo, showing no batch effect across experiment.

**Supplementary Figure 4.**
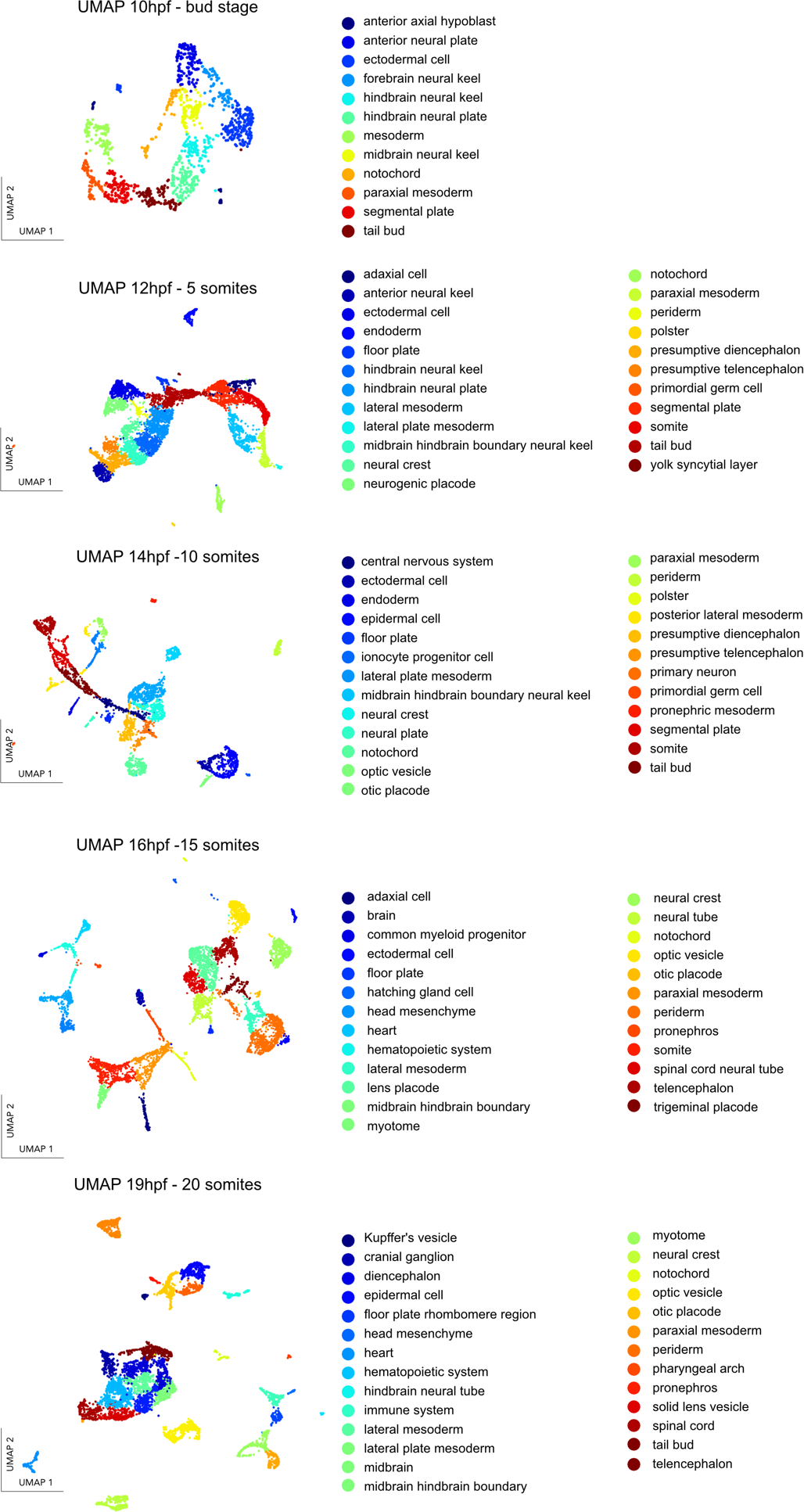
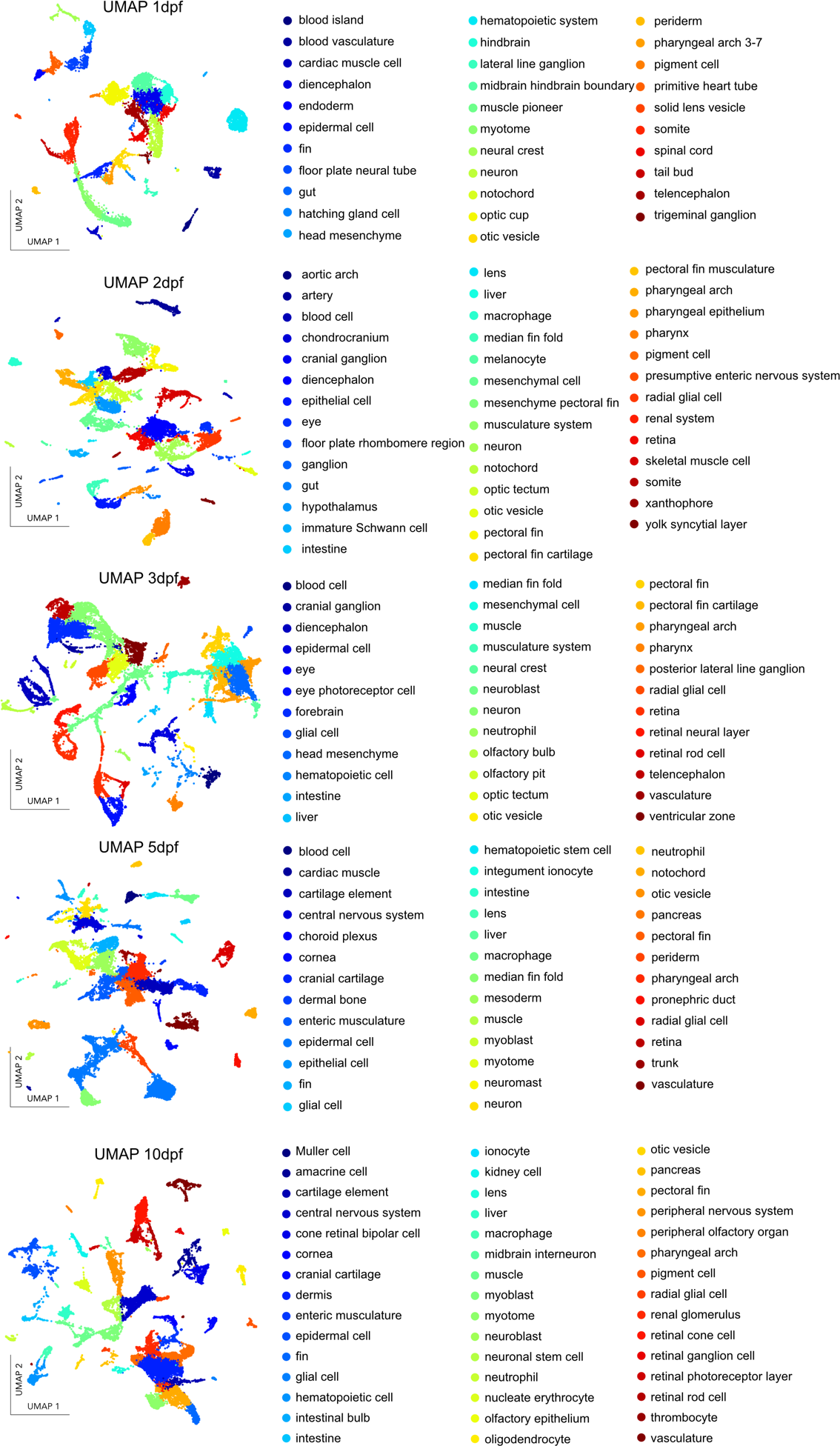
Summary for the cell type annotations represented on each timepoint UMAPs. We annotated 529 clusters based on gene enrichment cross-checked with published articles (see Methods).

**Supplementary Figure 5.**
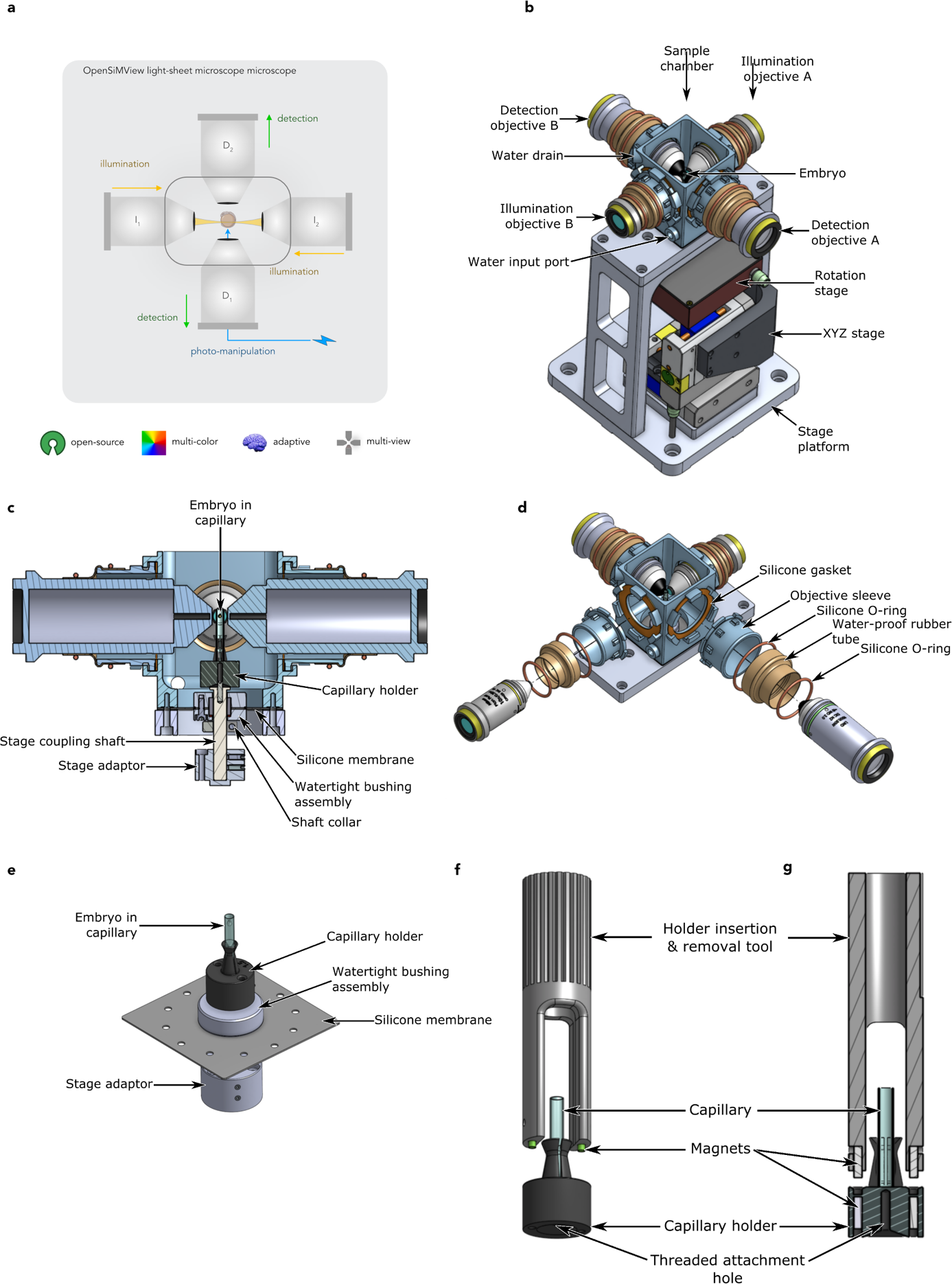
Structure of the sample chamber The sample chamber is designed to hold the sample (fish embryo) immersed in water and provide water-tight ports for two illumination and two detection objectives (allowing the objectives to move in 3 dimensions) aimed at the sample, and a water-tight connection to the combined XYZ and rotation (XYZR) stage that moves the sample with those four degrees of freedom. **a.** Overview of the multi-view light-sheet microscopes used in this study: OpenSiMView –two detections and two illuminations multi-view light-sheet microscope. **b.** The sample chamber is mounted on a custom-machined aluminum holder that places it above the XYZR stage, which is in turn mounted onto a custom-machined aluminum platform. The sample (fish embryo) sits in the middle of the chamber, surrounded by the four objectives. **c.** The sample chamber itself is composed of the following parts: (1) A main body (3D printed) with openings for the four objectives and input and output ports for water. (2) Four objective sleeves (3D printed) that connect to the chamber body using a twist-lock bayonet mechanism. (3) Laser-cut silicone gaskets that provide a water-tight seal between the objective sleeves and the chamber body. (4) Four rubber tubes that create a flexible and water-tight coupling between the objective sleeves and the objectives, while allowing movement of the objectives in all 3 dimensions relative to the sample. These tubes are made by cutting sections of the fingers from nitrile laboratory gloves. (5) Silicone O-rings that secure the rubber tubes to the objective sleeves and objectives, to reduce the chance of water leakages. (6) A custom-machined aluminum base to which the chamber body is bolted **d.** Cross section of the sample chamber, along the main axis of the detection objectives, showing the fish embryo in the capillary. The capillary is mounted onto a 3D printed capillary holder that thread onto a custom-machined stainless-steel shaft that couples to the XYZR stage. This shaft passes through a laser-cut silicone membrane clamped between the chamber body and the chamber base, via a water-tight bushing composed of two 3D printed halves that clamp onto the silicone membrane and hold in place a tubular silicone gasket (section of silicone tubing) that seals around the shaft, with silicone grease applied as lubricant between the tube and the shaft. The bushing assembly is designed such that the tubular silicone gasket is slightly compressed axially, to increase the sealing pressure against the shaft to avoid water leaks. The shaft can freely rotate inside the bushing (driven by the rotation stage), while the silicone membrane allows the shaft to move in 3 dimensions (driven by the XYZ stage). A 3D printed collar around the shaft limits the vertical sliding of the shaft in the bushing to about 2mm. The shaft is connected to the rotation stage via a custom-machined aluminum adaptor. **e.** Detail of the capillary holder showing the top half of the bushing assembly and the silicone membrane. The circular pattern of holes on the membrane are for the screws that attach the chamber body to the chamber base. **f.** Angle view. **F.** Cross section view of a 3D printed tool is used to easily insert the capillary holder into, and remove it from, the sample chamber. The tool has two 1/16” (1.587mm) diameter neodymium magnets (axially magnetized) that mate into corresponding cavities in the capillary holder. Identical magnets are embedded into the capillary holder (right underneath the tool-mating cavities) to provide a mating force between the tool and the holder. All magnets are glued in place using epoxy adhesive. With this design the tool can transfer the capillary holder to the chamber and screw it onto the stage-coupling shaft (or remove it from the shaft). The magnetic force produced by the magnets is strong enough to hold the weight of the capillary holder, and the magnet-to-cavity coupling allows the transfer of torque for screwing the holder onto the shaft. All 3D printed chamber parts and holder were printed on Formlabs Form 2 and Form 3 3D printers using Formlabs Tough v5 resin.

**Supplementary Figure 6.**
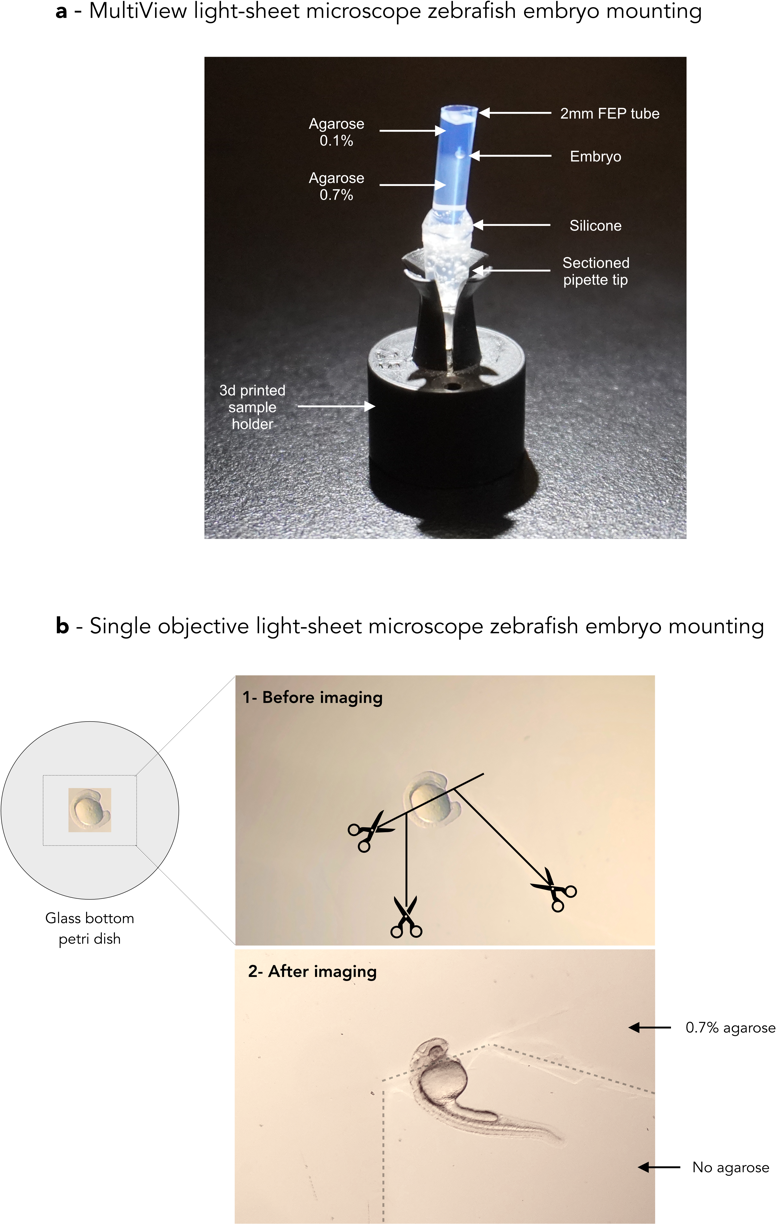
Mounting strategies for long-term zebrafish imaging in the light-sheet microscope. a. For the multiview light-sheet microscope the zebrafish are mounted in FEP tubes in 0.1% low melting point agarose in embryo medium supported by a 0.7% % low melting point agarose (Sigma, A0701). The low concentration agarose allows the full embryo development with tail elongation. The FEP is sealed in a pipette tip sealed with low-toxicity silicone (Kwik-Sil^tm^ from World Precision Instrument) and then placed in the capillary holder described Fig. S5 and placed in the chamber using the capillary holder insertion & removal tool. b. Embryo is placed in a 35mm cell culture dish with a 20mm glass bottom (Stellar Scientific) in 0.7% low melting point agarose (Sigma, A0701). After correct positioning the agarose surrounding the posterior part is removed, to not prevent tail extension.

**Supplementary Figure 7.**
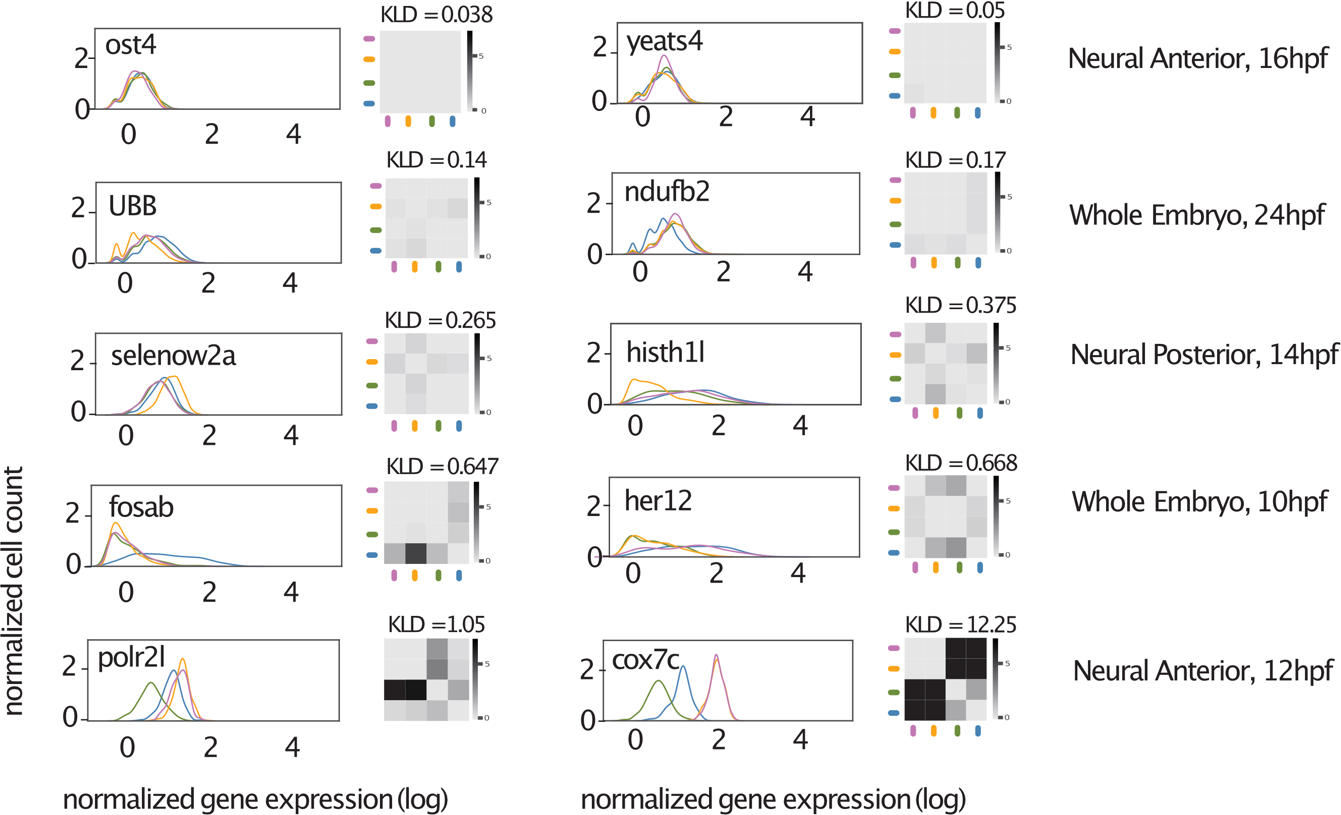
Examples of genes with different KLD scores. Distributions of log normalized gene expression grouped by embryo for genes with increasing values of KLD (0.038:12.25). The heatmaps show KL(x,y) across all pairs of embryos (embryos compared to themselves along the diagonal have KL = 0). Each gene’s KLD score (the median of the pairwise KL values, excluding the diagonal) is shown above the heatmap.

**Supplementary Figure 8.**
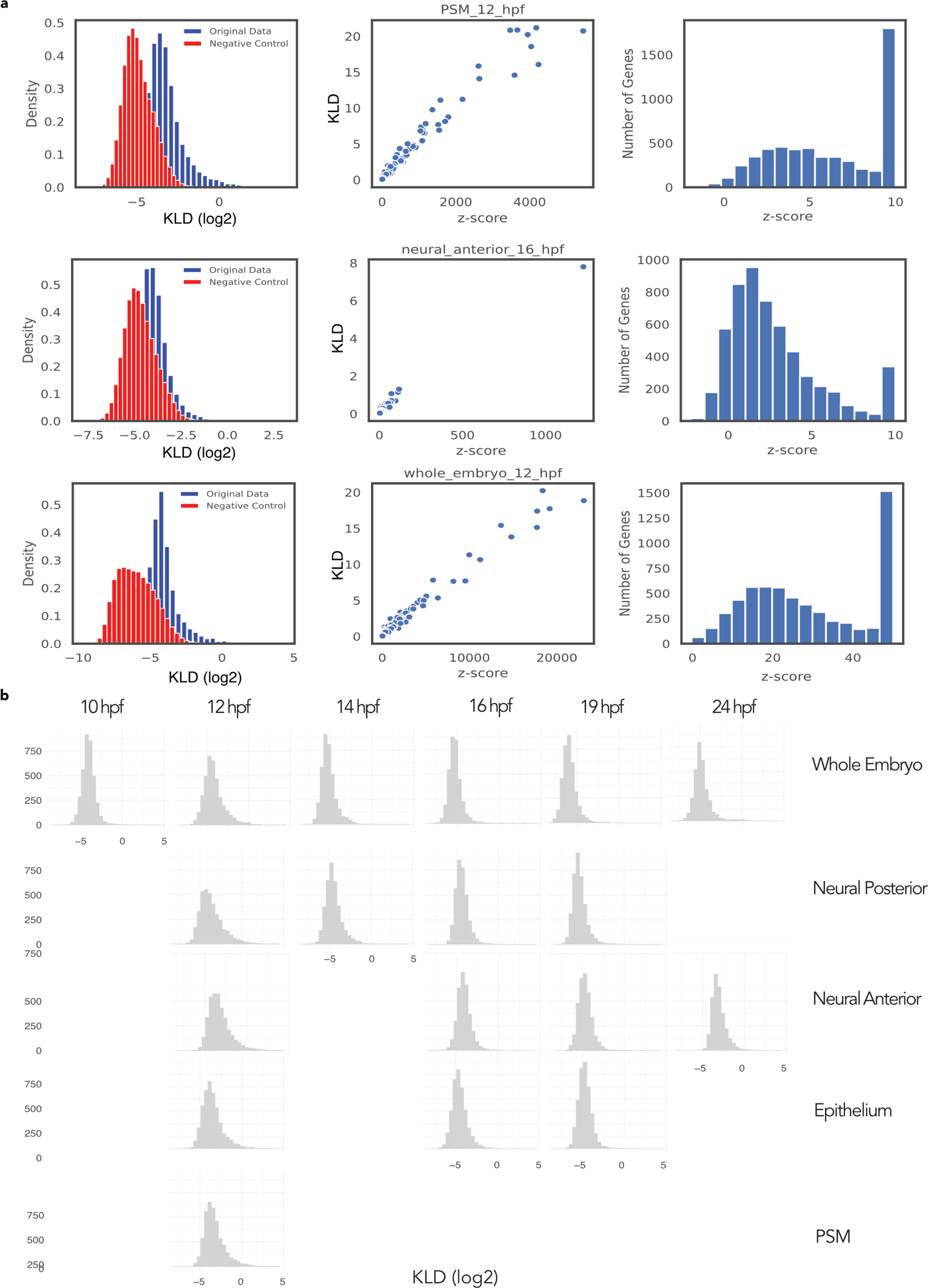
KLD score control and distribution during development (continued next page). a. Negative control for the KLD scores. We generated a null distribution for the KLD scores by randomly reshuffling the embryo ID of individual cells. The null distribution allows testing the hypothesis that the differences observed between the four embryos can be attributed to random samples from the same distribution. For each gene, we randomized the embryo IDs of individual cells, re-calculated the KLD scores (red histograms), and compared them to the original distributions (blue histograms), repeating the randomization and calculation process 20 times. We calculated the null distributions for different cell types, including the whole embryo. The scatterplots show the z-scores, computed for each gene as the difference between the null distribution (20 null KLD scores for each gene) and their true KLD score. We found a strong positive correlation between the z-score and the KLD score and decided to use the latter. The histograms of z-scores show the fraction of genes with significant KLD (z-score >10). We show the null distributions and z-scores for the whole embryo (12hpf), the presomitic mesoderm (PSM, 12hpf), and the Neural anterior cells (16hpf) as representative examples. b. KLD distributions across all genes for different cell types and time points. The histogram shows the KLD scores for all genes that passed the quality filters (fraction of cells with zero counts < 0.8, mean log-normalized expression > 0.2, at least 80 cells per fish for each cell type/timepoint) for that subset of cells. Missing histograms correspond to cell types that don’t have enough cells per embryo (minimum of 80).

**Supplementary Figure 9.**
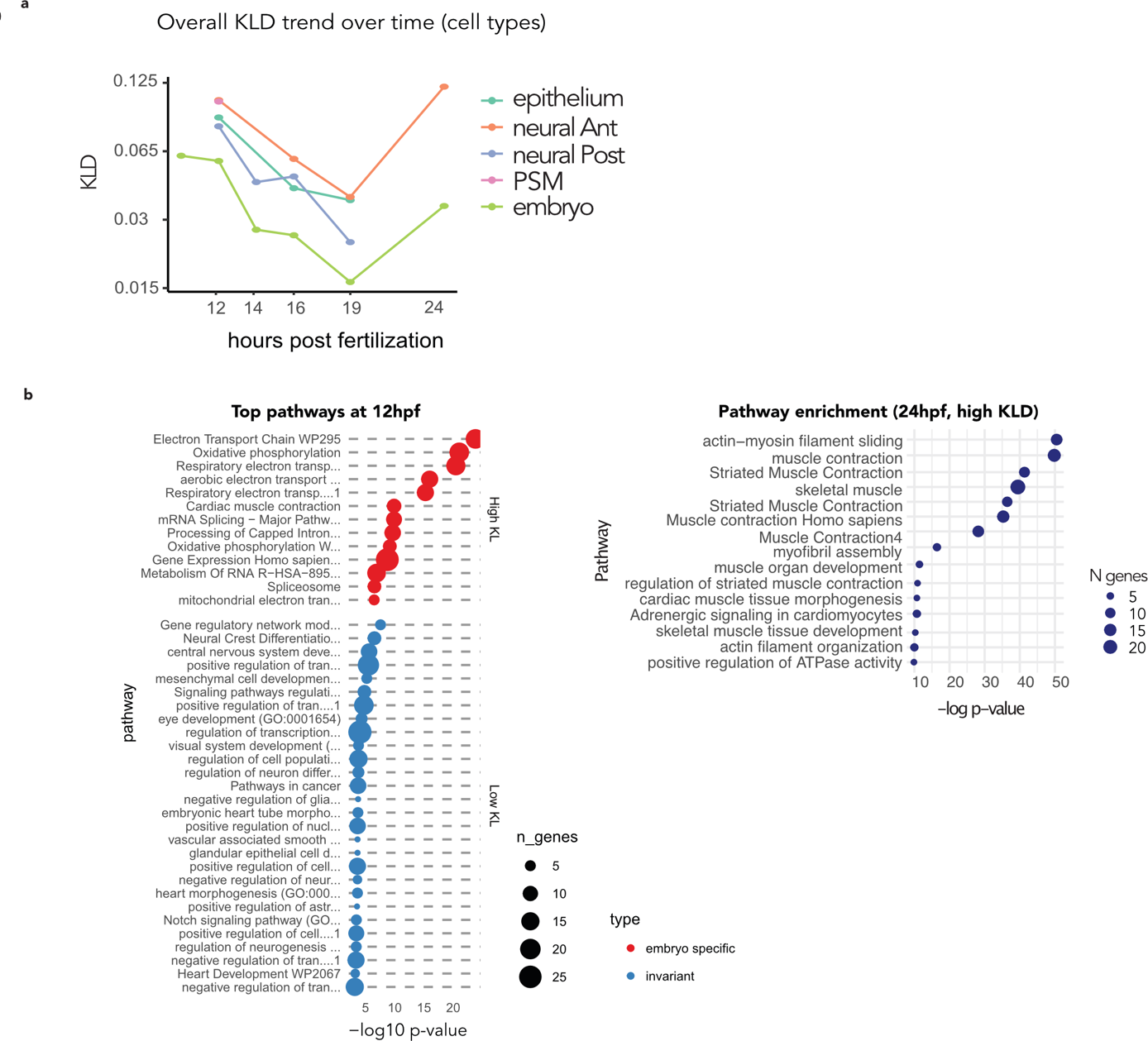
KLD and enrichment pathway details overtime. a. KLD trends for different cell types. The KLD median value across all genes for different cell types across time points. Each dot represents the median across all genes that passed the quality filters for that cell type. Some cell types do not have enough cells at some time points and therefore do not appear in the plot. b. Top pathways at 12 and top pathways at 24. An extended list of enriched pathways in high and low KLD genes for the whole embryo at 12hpf. For 24hpf, we show the top enriched pathways in high KLD genes.

**Supplementary Figure 10.**
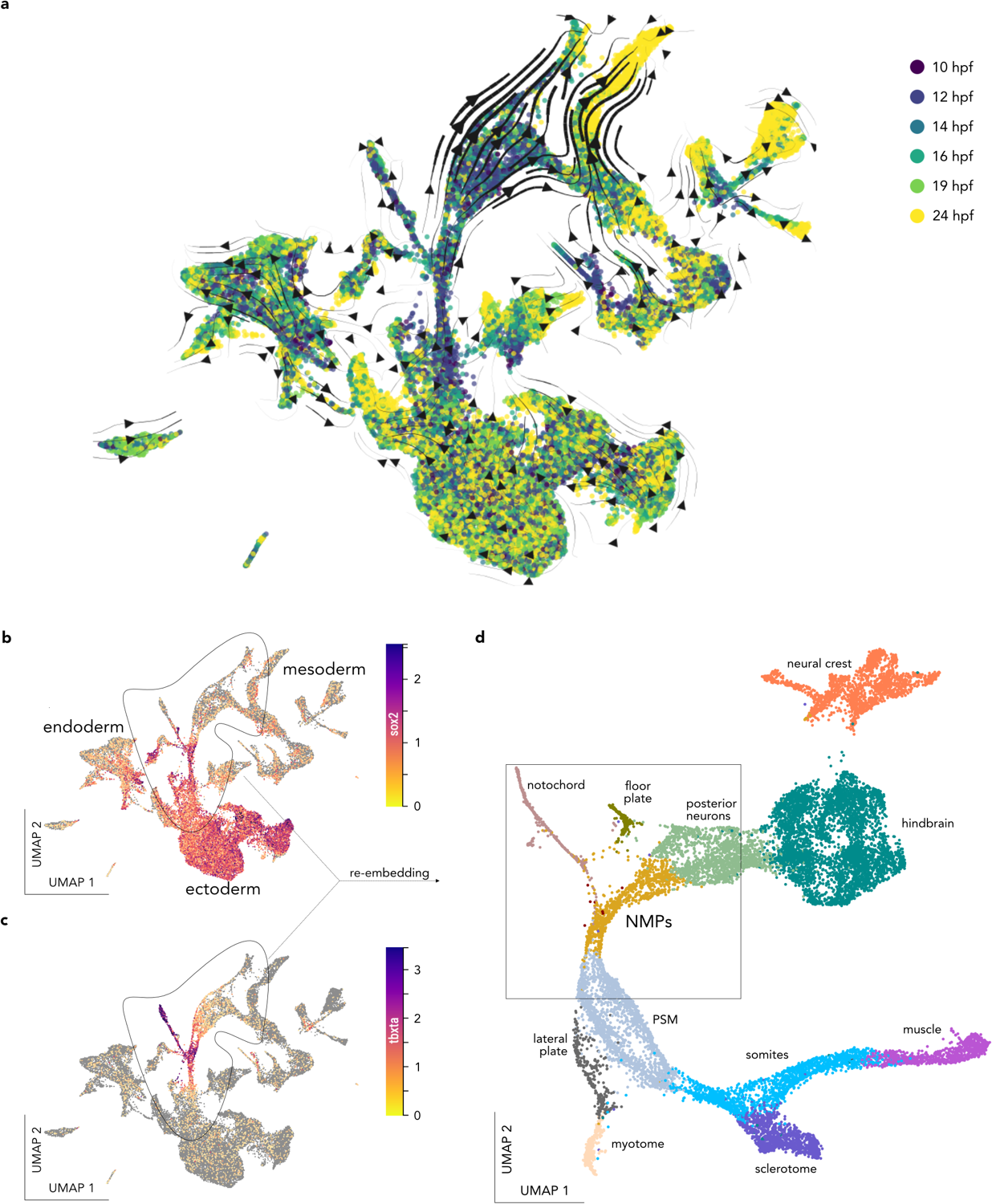
Early time points UMAPs (10 hpf, 12 hpf, 14 hpf, 16 hpf, 19 hpf and 24hpf) and the NMP cluster. a. UMAP of early zebrafish development (integration of 10 to 24 hpf time points in a single object), color codes by time points, the arrow streams correspond to the RNA velocities. b. Early time points UMAP embedding color coded for sox2. c. Early time points UMAP embedding color coded for tbxta. d. The cluster co-expressing sox2 and tbxta (molecular definitions of the NMP cells) and there neighboring clusters are re embedded in a new UMAP.

**Supplementary Figure 11.**
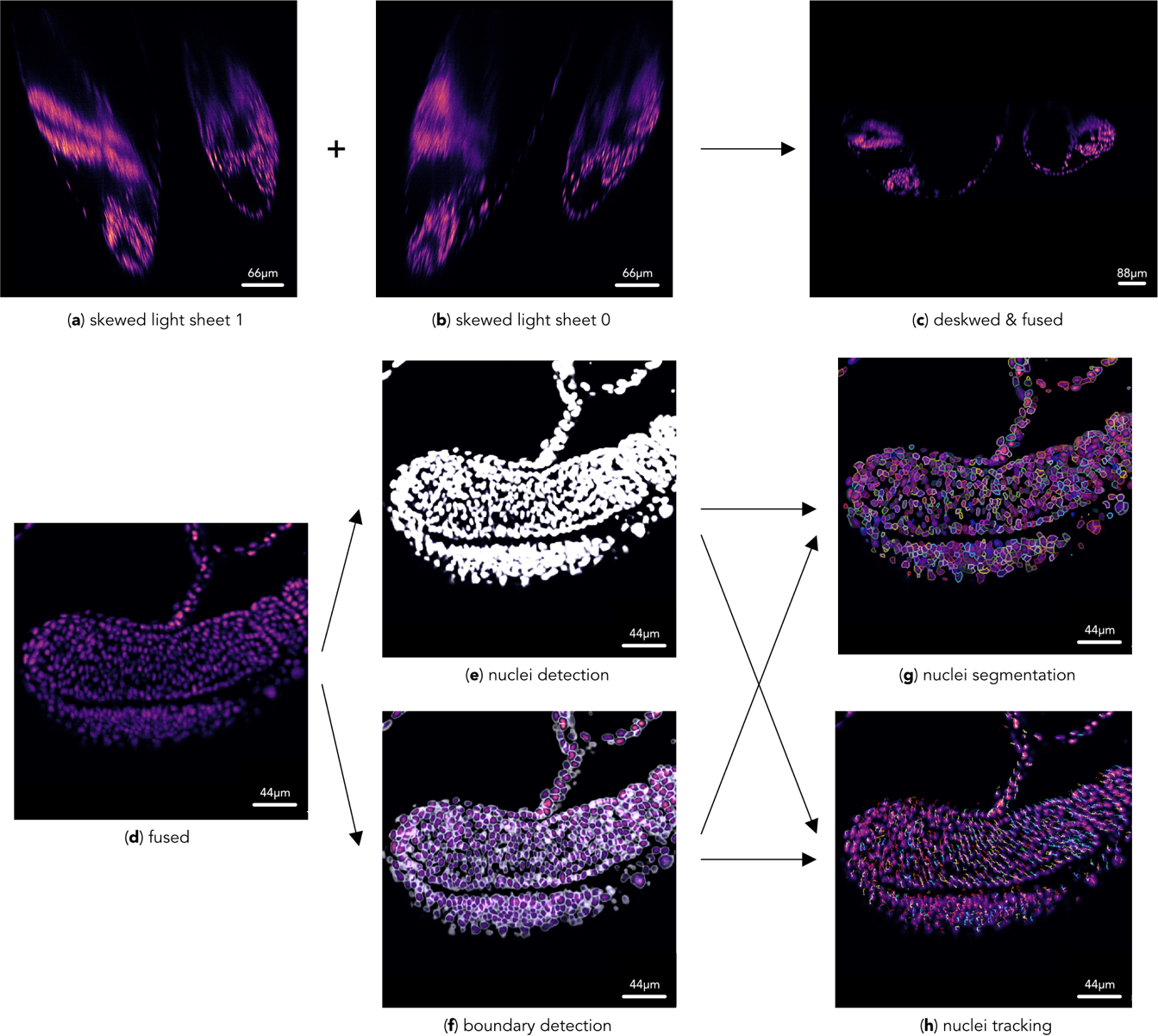
Single-objective light-sheet microscope image processing pipeline. a. Maximum intensity projection skewed views from light sheets 1 and 2; b. Same as a. c. Maximum intensity projection of deskewed and fused volume. d. xy-slice of embryo’s tail. e. Neural network cell detection of region. f. (Neural network cell boundary prediction of region (d). (g, h) Results from joint segmentation and tracking. g. Segmentation masks. h. Respective tracks.

**Supplementary Figure 12.**
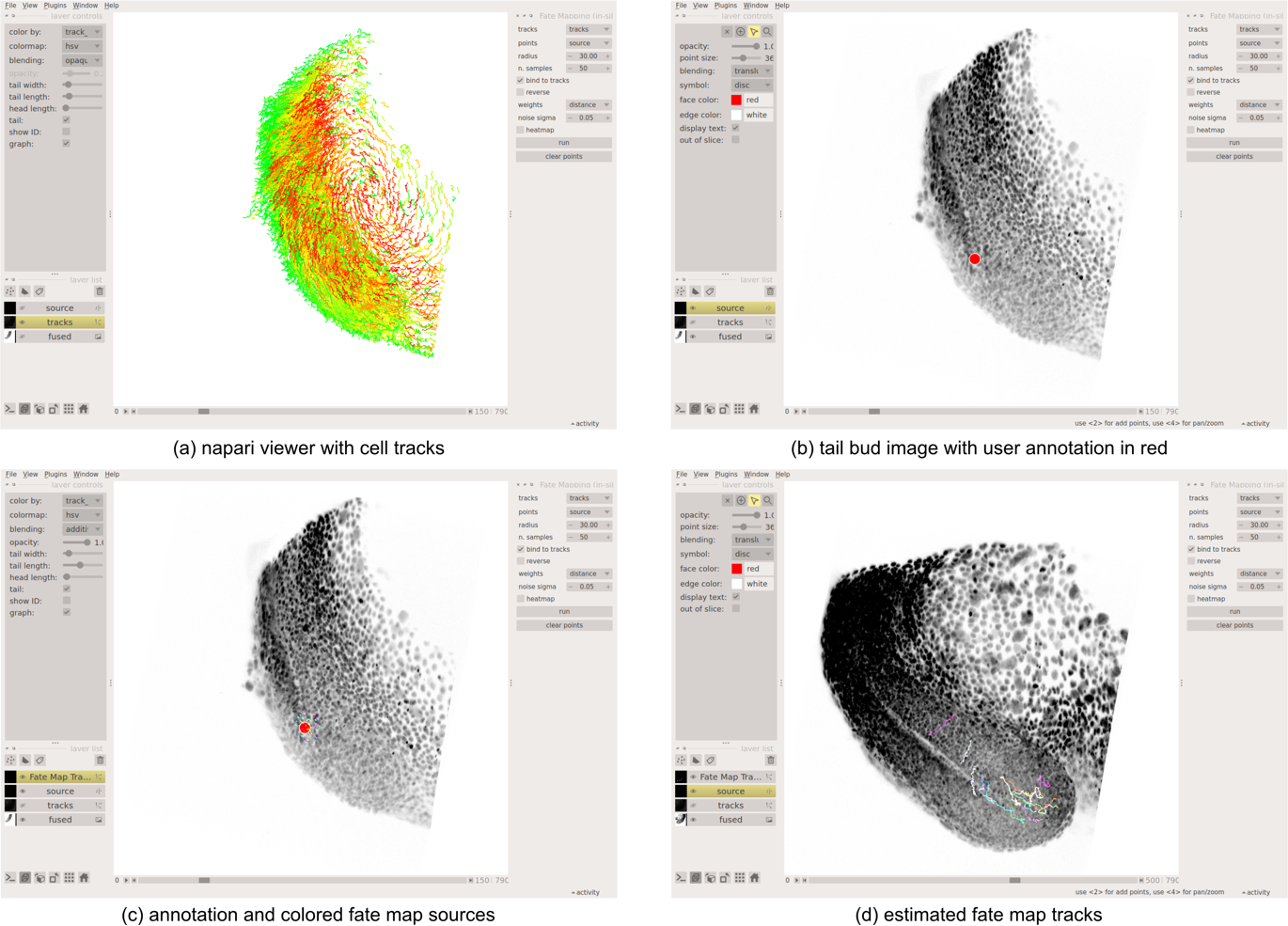
The *in silico* tracking plugin for Napari. a. Visualization of cell tracks in Napari. b. Selection of region of interest. c. Propagation of stepwise local movement estimation. d. Display of the estimated fate map tracks.

**Supplementary Figure 13.**
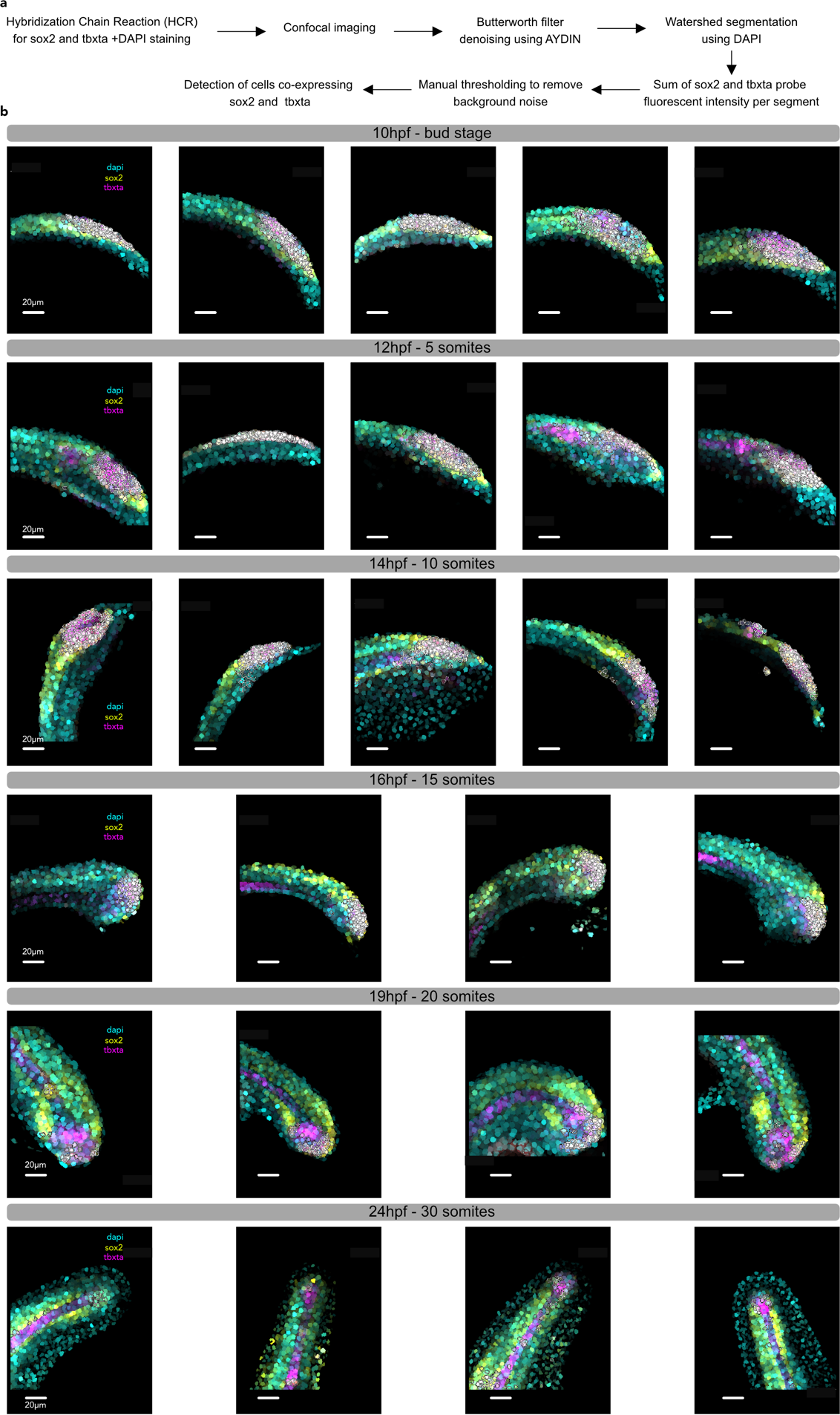
Spatial identification of NMPs presumptive territory using Hybridization Chain Reaction (HCR) during somitogenesis (continued next page). a. The pipeline used to identify the NMP, image the zebrafish tail, segment the DAPI channel, and finally identified the NMP (co-expressing *sox2* and *tbxta*). b. Processed images showing the evolution over time of the cells co-expressing *sox2* and *tbxta* - NMP presumptive territory.

**Supplementary Figure 14.**
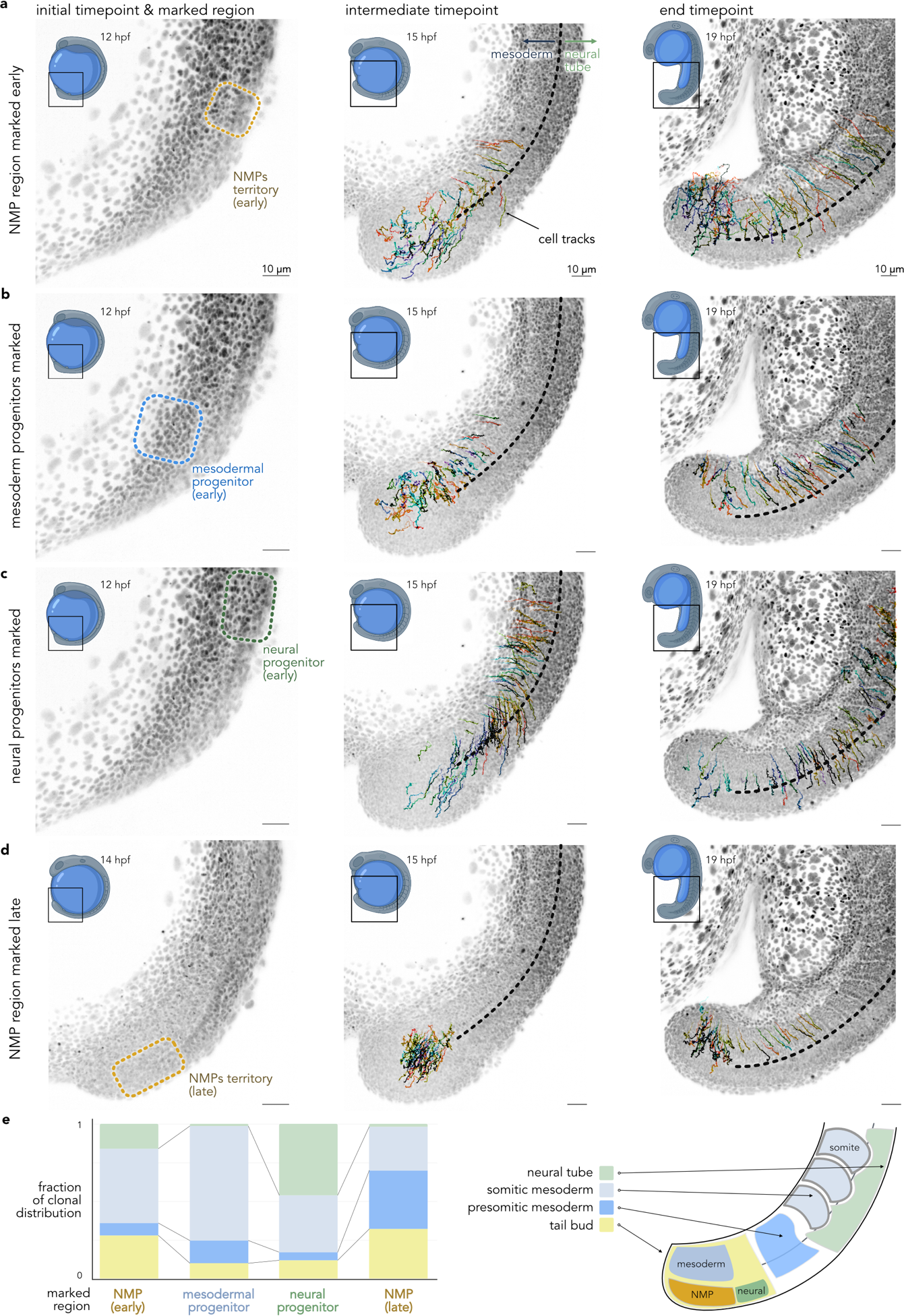
Reconstruction of the zebrafish embryonic posterior progenitor’s territory via *in silico* fate mapping. **a.** *In silico* fate mapping of the pluripotent early NMP territory. Left - selected territory at the initial timepoint, middle - intermediate timepoint showing tracks for selected cells, right – final time point. **b.** *In silico* fate mapping of the axial mesodermal posterior progenitors. We observe that these cells mostly integrate into the mesoderm (74%). **c.** *In silico* fate mapping of the axial neural posterior progenitors. These cells partly integrate the neural tube (46%). **d.** *In silico* fate mapping of the late presumptive NMP territory. These cells integrate mostly the mesodermal tissue (70%). **e.** Left – the proportion of the clonal distribution after photoconversion of the different progenitor territories: NMPs early (total number of cells = 75), mesodermal progenitors (total number of cells = 81), neural progenitors (total number of cells = 76), NMPs late (total number of cells = 53). Right – a schematic representation of the different axial progenitor territories.

**Supplementary Figure 15.**
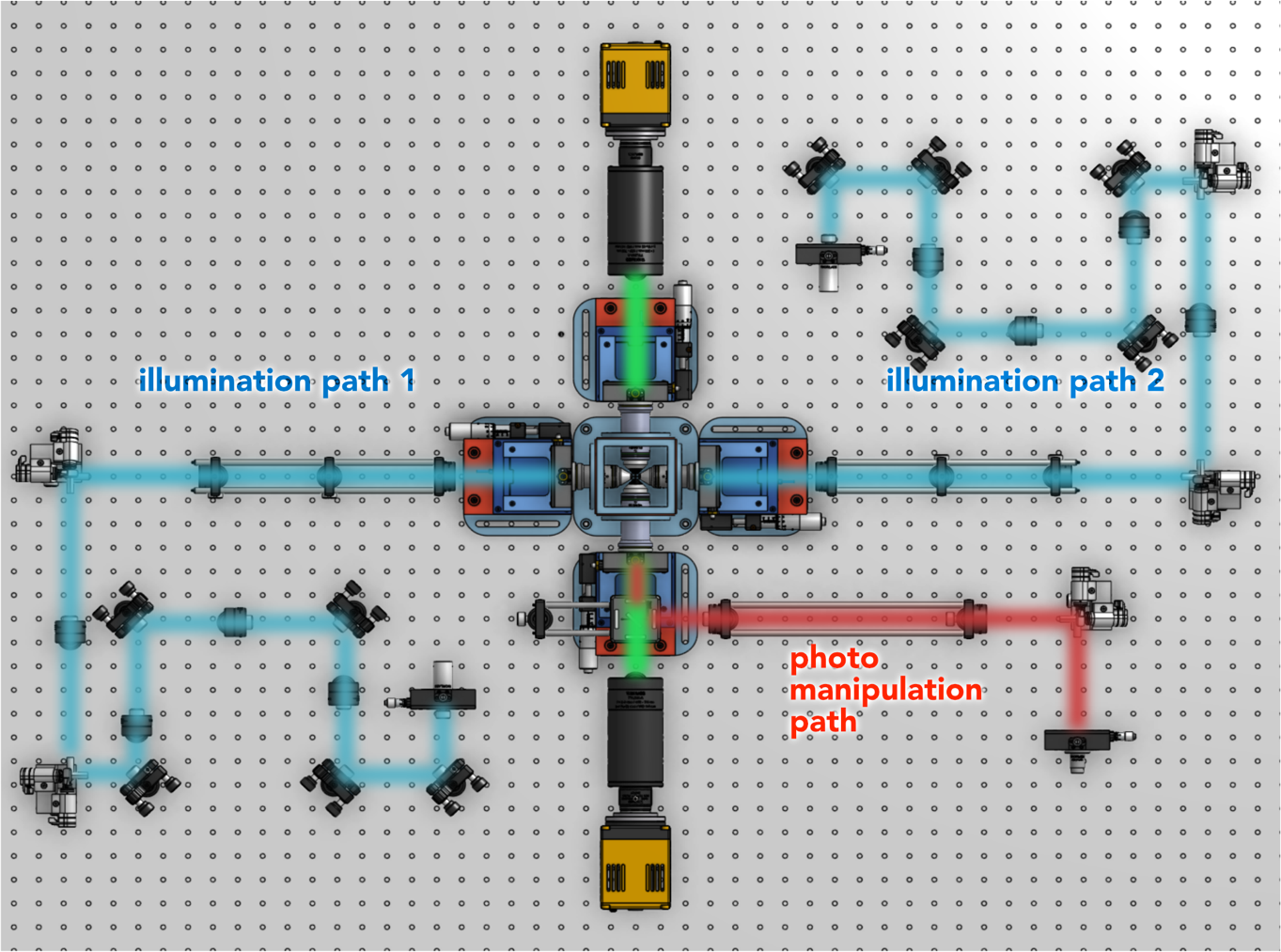
Illustration of the illumination paths (in blue) and photoconversion arm (in red), in our multiview light-sheet microscope.

**Supplementary Figure 16.**
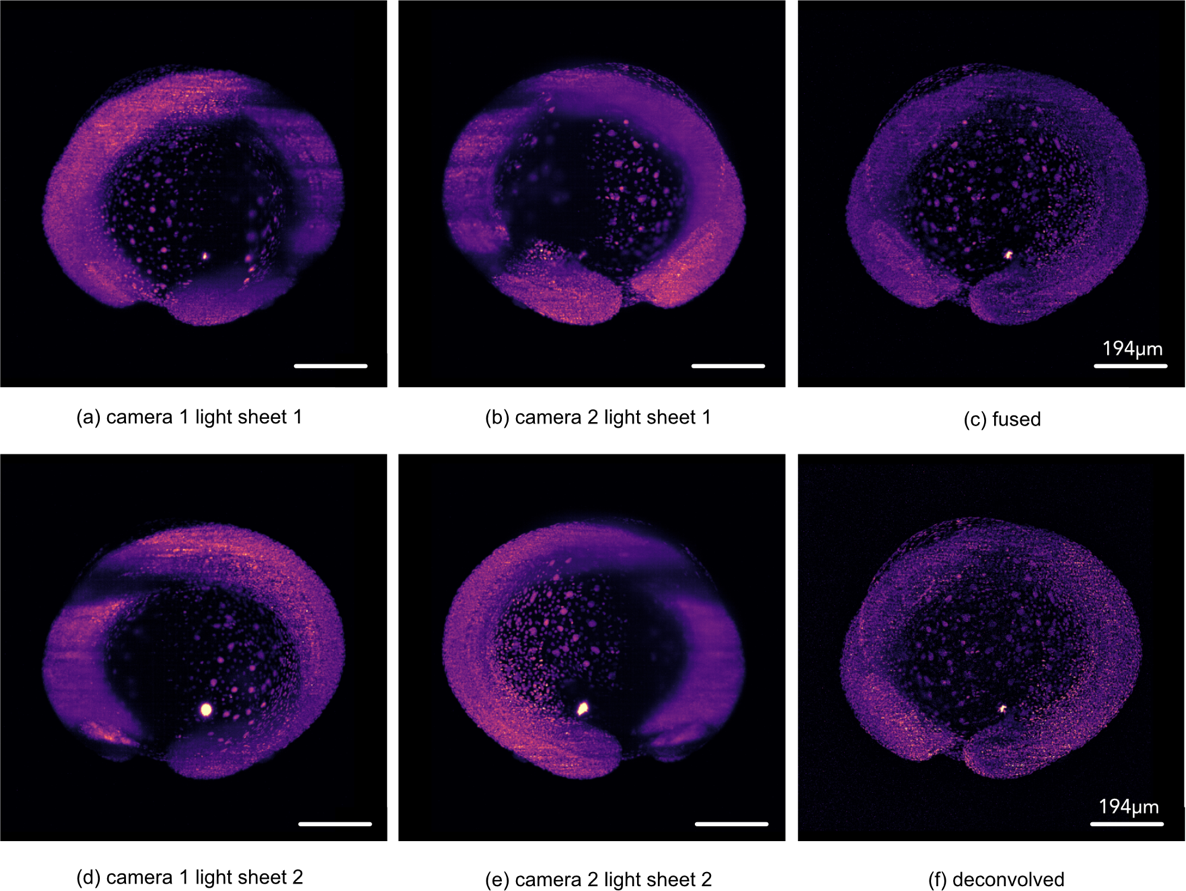
Intermediate results of multi-view light-sheet microscope image processing pipeline. Volumes shown as maximum intensity projection on the z-axis of: a. Camera 1 light sheet 1; b. Camera 2 light-sheet 1; c. Camera 1 light sheet 2; d. Camera 2 light sheet 2; e. Fused, camera 2 volumes are flipped before fusion. f. Deconvolved volume after fusion.

**Figure 17.**
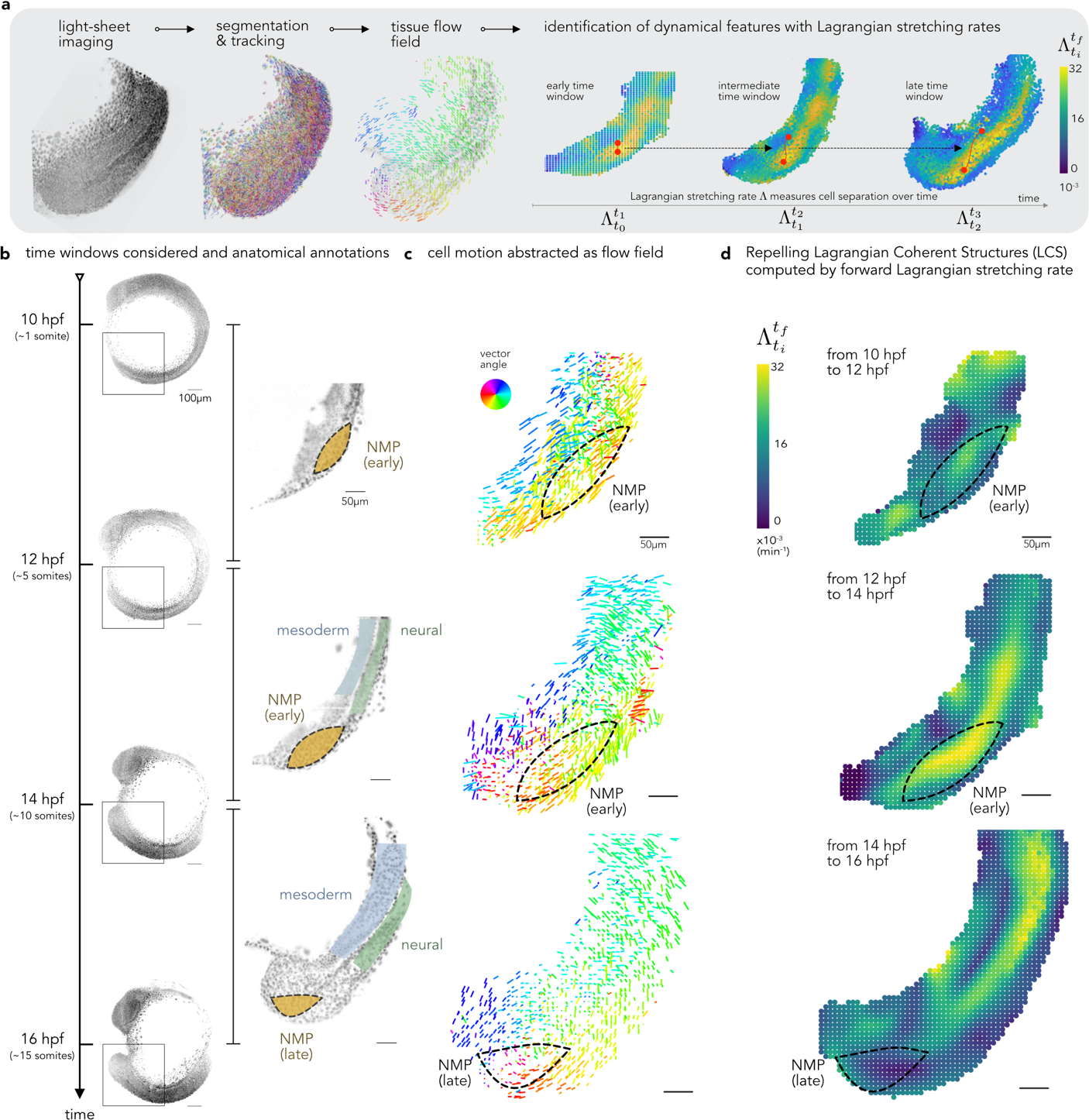
Tissue morphodynamics constrain the state transition of NMPs during vertebrate body axis elongation. **a.** Sketch of the dynamic morphoskeleton approach used to identify robust dynamical features from multicellular flows. We focus on repellers – a type of Lagrangian coherent structure locating divergent cell trajectories. We identify repellers by computing the Lagrangian stretching rate (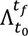, see Supp. Theory Note). **b.** Illustration of the time intervals selected to compute the stretching rates, and anatomical annotations of the tail’s territories (NMP, neural and mesodermal tissues). **c.** Vector fields that represent the cells’ motion during these time intervals, and dashed lines represent the presumptive NMP territory. **d.** Lagrangian stretching rate (brighter shades correspond to more stretching i.e., divergent cell trajectories) and the emergence of repellers in the early NMP presumptive territory. This territory was ascertained by HCR and is outlined by a dashed line.

**Supplementary Figure 18.**
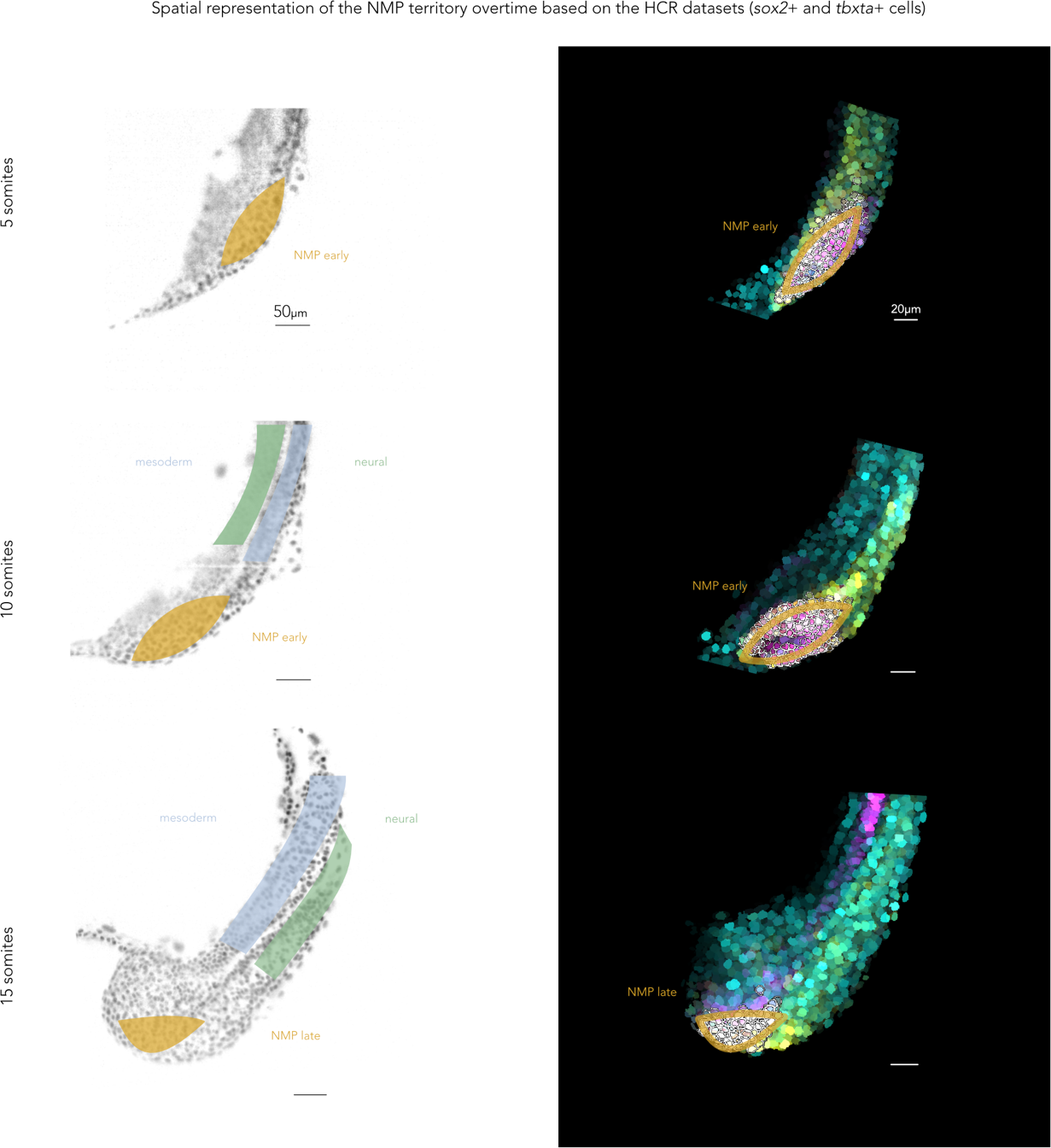
Spatial representation of the NMP presumptive territory used for the FTLE analysis is based on the HCR staining (cell double positive for *sox2* and *tbxta*), at 5somites or 12hpf, 10somites or 14hpf and 15somites or 16hpf.

**Supplementary Figure 19.**
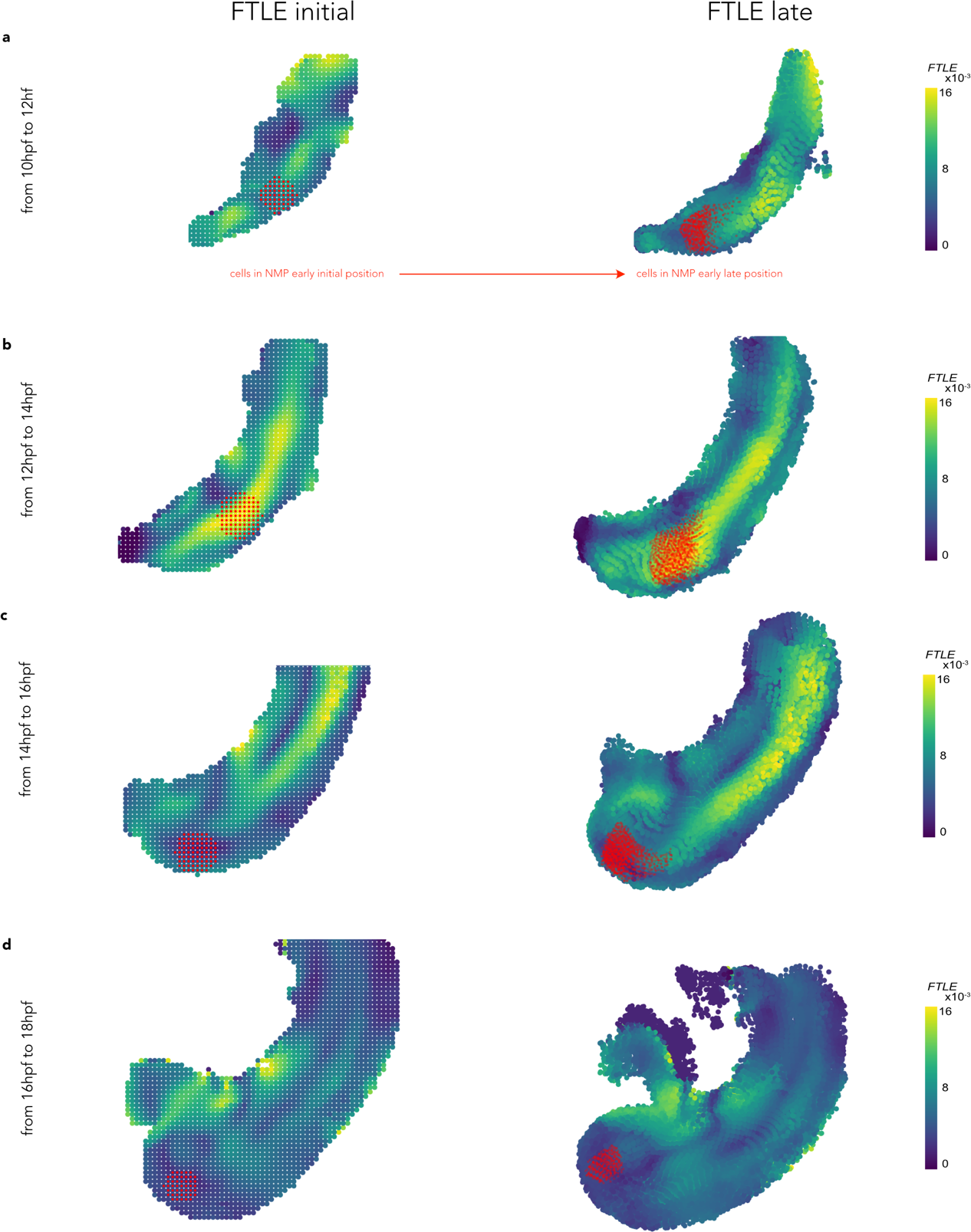
Temporal dispersion of putative NMP cells when computing the forward FTLE in Lagrangian Coherent Structures for different temporal windows. Labelled putative cells are represented in red in their initial location (FTLE initial) and their destination (FTLE late). In NMP early progeny is distributed in both mesodermal and neural tissue. In late NMP, the progeny only integrates the mesoderm.

**Supplementary Figure 20.**
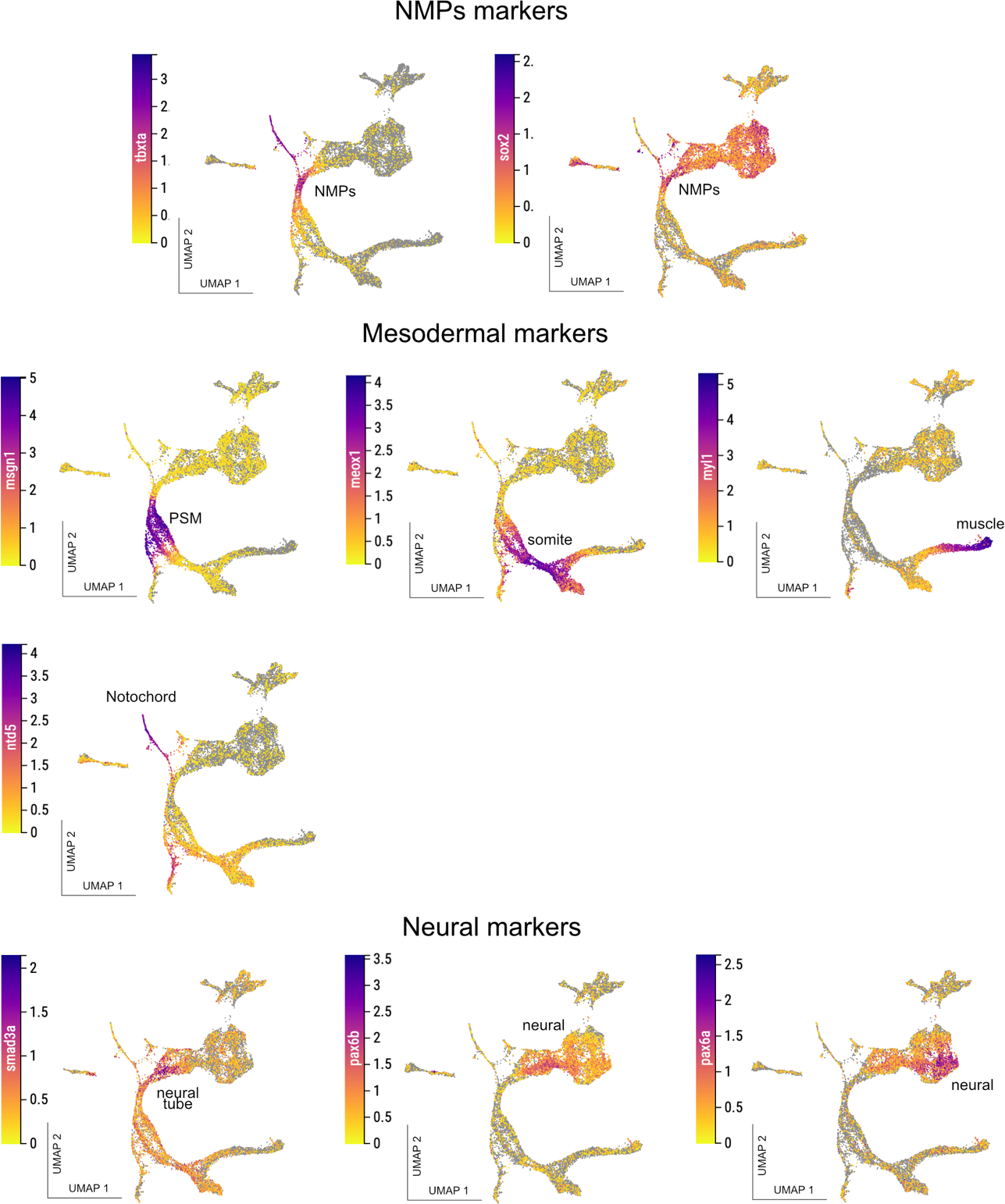
NMP cells UMAP embedding color-coded for top genes important for the NMPs (top), mesodermal differentiation (middle) and neural differentiation markers (bottom).

**Supplementary Figure 21.**
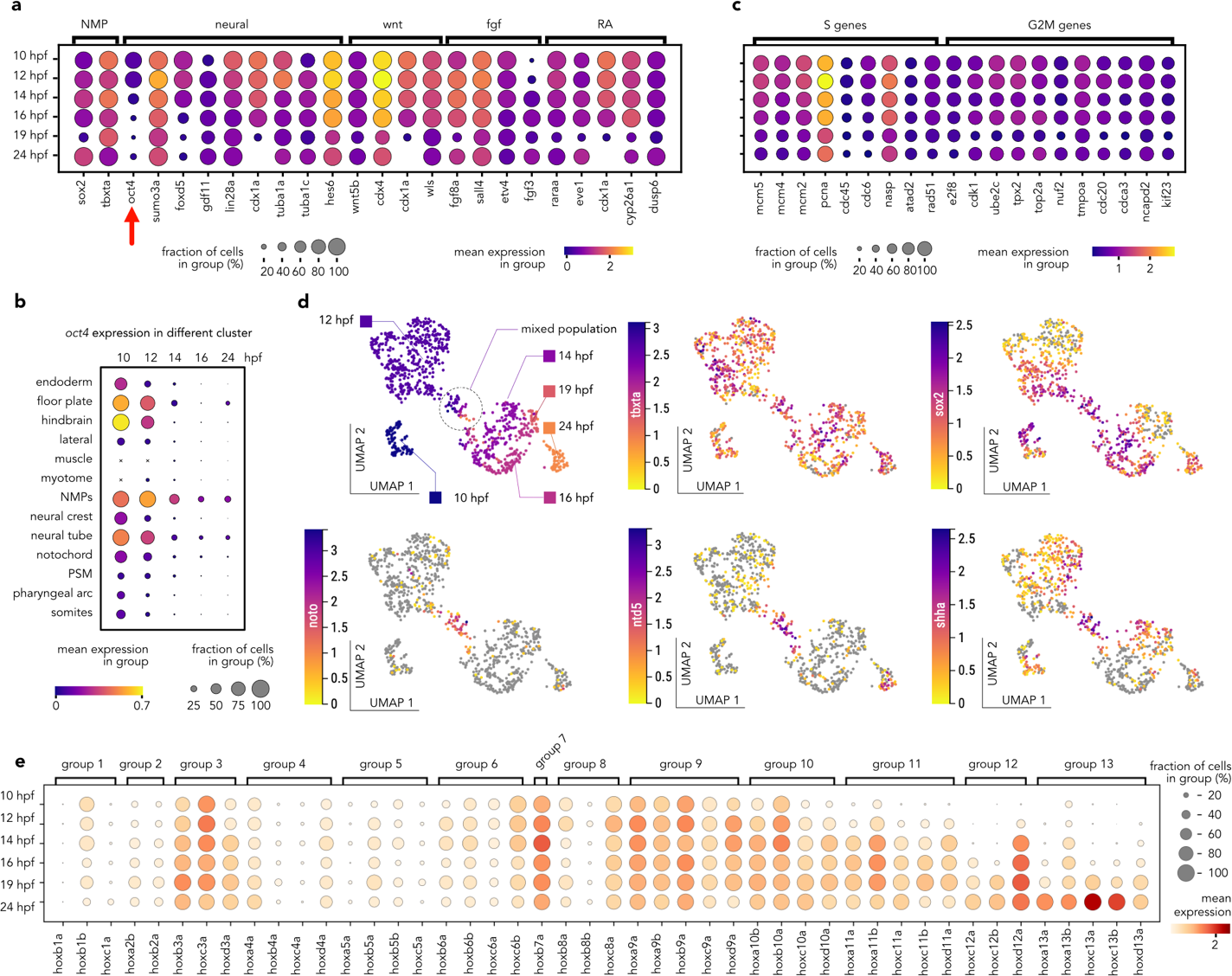
NMP cells molecular fingerprints extracted from the scRNAseq a. Selection of NMPs enriched genes for signaling pathway important for vertebrates’ axial elongation showing a peak of expression early during development. b. Selection of important genes for cell cycle enriched in the early NMPs. c. Dotplot of the zebrafish hox genes expression overtime in the NMP clusters, showing a collinear activation of hox pathway. d. Zebrafish oct4 (pou5f3) timeline of spatial expression in the posterior body clusters, showing late maintenance of oct4 specifically in the NMP cells. e. UMAP embedding of the NMPs cluster in the early integrated UMAP (10hpf, 12hpf, 14hpf, 16hpf, 19hpf, and 24hpf) color-coded by time point (top left). The same embedding is then color coded for the NMP molecular signature genes (sox2 and tbxta) and then for three genes very important for notochord development.

## SUPPLEMENTARY THEORY NOTE

Any framework to analyze spatio-temporal trajectories in morphogenesis requires a self-consistent description of cell motion that is independent of the choice of reference frame. This frame-invariant property, called objectivity^1^, is a fundamental axiom of mechanics and ensures that the description of deforming biological tissue is independent of the coordinate frames we choose to describe its motion. Non-objective metrics, for instance, will yield different (inconsistent) results if cell velocities are described from a frame moving with a drifting embryo or a fixed lab frame (see e.g., Figure 1 in the reference^2^). The velocity field and the streamlines, which are trajectories of the frozen velocity field, are common nonobjective metrics^1^.

To analyze cellular flows generated in the zebrafish tailbud, we use the recently developed Dynamic morphoskeletons (DM)^2^, which is based on the notion of Lagrangian Coherent Structures (LCS)^3^. The DM is objective and founded on a Lagrangian description of tissue deformation captured by Finite Time Lyapunov Exponents (FTLE), which naturally combine local and global mechanisms along cell trajectories. Given a time interval *T* = *t_f_* – *t_0_* of interest, the DM locates repellers from which cells diverge, attractors towards which cells converge, and their domain of attraction identifying the initial positions of the cells that will end up in the attractor. The DM reveals the intrinsic geometric feature of spatiotemporal trajectories and is robust to noise^2^, hence it is ideal for quantifying morphogenesis.

Since morphogenetic flows do not undergo exponential stretching, here we identify the DM by computing the Lagrangian stretching rate, a simpler but equivalent version of the FTLE. Because we are specifically interested in the compartmentalization of the NMP region, we compute the forward Lagrangian stretching 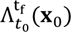 to identify the repelling spatiotemporal regions (repelling LCSs) in the embryo. Repellers are marked by high values of 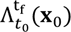. The Lagrangian stretching rate field 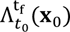 computed for the time interval and initial point ***x***_0_ is given by,

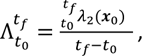

where 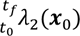 denotes the largest singular value of the Jacobian of the flow map 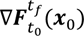 and 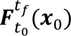 is the flow map describing the cell trajectories from their initial position ***x***_0_ to the final position 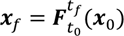. We compute 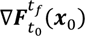 from cell trajectories using a recently developed technique to identify LCSs from sparse and noisy trajectories^4^. This method relies on calculating the separation between neighboring trajectories over time. For an initial cell position located at ***x***_0_, neighboring trajectories are those starting within a sphere of radius *δ* from the point ***x***_0_. For all the results shown in this paper, *δ* = 100. We use the publicly available code in the reference^4^ and their strategy to select the optimal *δ* from data. The Lagrangian stretching 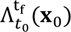 describes a deformation rate, it has units of 1/time, and it quantifies the maximum Lagrangian deformation rate over the time interval [$_#_, $_"_] of a region whose initial position is centered at ***x***_0_ (Figure 1a). Nearby cells on the opposite sides of a repelling LCS are maximally separated (Figure 1b), facilitating cell populations to achieve differential mechanical environments for cell fate specification.

**Fig 1:**
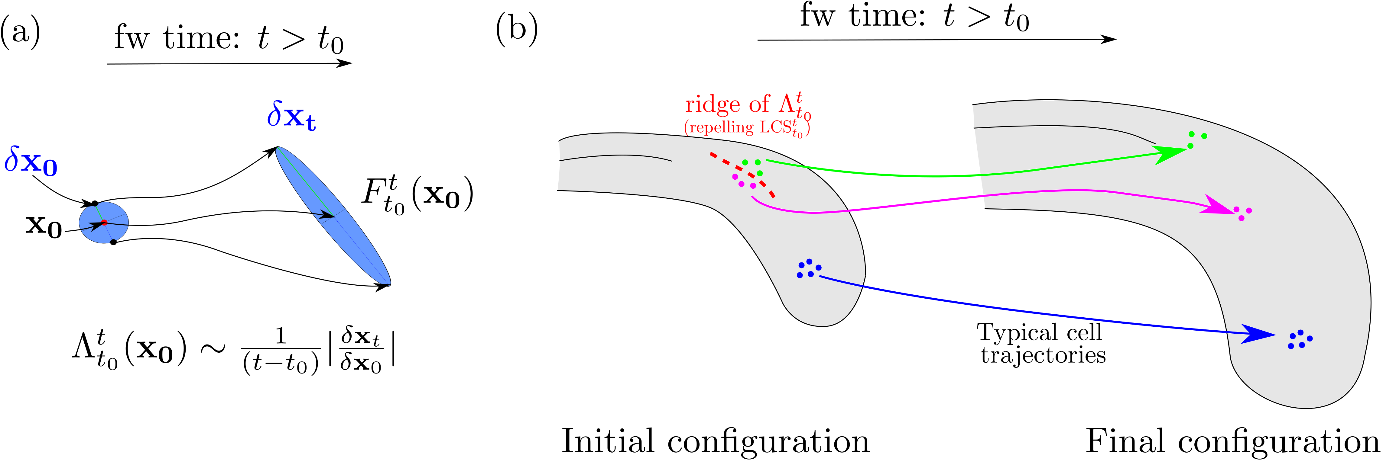
(a) The Lagrangian Stretching 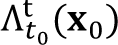 is a measure of the maximum separation generated by the flow over the time interval [*t*_0_*t*] between two nearby points in the neighborhood of ***x***_0_. (b) Ridges (ie sets of high values) of 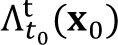 mark repelling LCS over [*t*_0_, *t*], which are the set of initial cell locations ***x***_0_ where the Lagrangian stretching rate is maximum.

## Dynamic Morphoskeletons quantify morphogenetic flows

Any framework to analyze spatio-temporal trajectories in morphogenesis requires a self-consistent description of cell motion that is independent of the choice of reference frame. This frame-invariant property, called objectivity^1^, is a fundamental axiom of mechanics and ensures that the description of deforming biological tissue is independent of the coordinate frames we choose to describe its motion. Non-objective metrics, for instance, will yield different (inconsistent) results if cell velocities are described from a frame moving with a drifting embryo or a fixed lab frame (see e.g., Figure 1 in the reference^2^). The velocity field and the streamlines, which are trajectories of the frozen velocity field, are common nonobjective metrics 1.

To analyze cellular flows generated in the zebrafish tailbud, we use the recently developed Dynamic morphoskeletons (DM)2, which is based on the notion of Lagrangian Coherent Structures (LCS)^3^. The DM is objective and founded on a Lagrangian description of tissue deformation captured by Finite Time Lyapunov Exponents (FTLE), which naturally combine local and global mechanisms along cell trajectories. Given a time interval *T* = *t_f_* – *t_0_* of interest, the DM locates repellers from which cells diverge, attractors towards which cells converge, and their domain of attraction identifying the initial positions of the cells that will end up in the attractor. The DM reveals the intrinsic geometric feature of spatiotemporal trajectories and is robust to noise ^2^, hence it is ideal for quantifying morphogenesis.

Since morphogenetic flows do not undergo exponential stretching, here we identify the DM by computing the Lagrangian stretching, a simpler but equivalent version of the FTLE. Because we are specifically interested in the compartmentalization of the NMP region, we compute the forward Lagrangian stretching 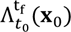 to identify the repelling spatiotemporal regions (repelling LCSs) in the embryo. Repellers are marked by high values of 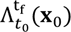. The Lagrangian stretching field 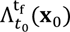 computed for the time interval [*t_0_*, *t_f_*] and initial point ***x***_0_ is given by,

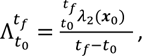

where 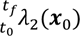 denotes the largest singular value of the Jacobian of the flow map 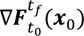 and 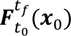 is the flow map describing the cell trajectories from their initial position ***x***_0_ to the final position 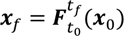. We compute 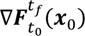 from cell trajectories using a recently developed technique to identify LCSs from sparse and noisy trajectories^4^. This method relies on calculating the separation between neighboring trajectories over time. For an initial cell position located a ***x***_0_, neighboring trajectories are those starting within a sphere of radius *δ* from the point ***x***_0_. For all the results shown in this paper, *δ* = 100. We use the publicly available code in the reference^4^ and their strategy to select the optimal *δ* from data. The Lagrangian stretching 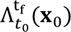 describes a deformation rate, it has units of 1/time, and it quantifies the maximum Lagrangian deformation rate over the time nterval [*t_0_*, *t_f_*] of a region whose initial position is centered at ***x***_0_ (Figure 1a). Nearby cells on the opposite sides of a repelling LCS are maximally separated (Figure 1b), facilitating cell populations to achieve differential mechanical environments for cell fate specification.

**Fig 1:**
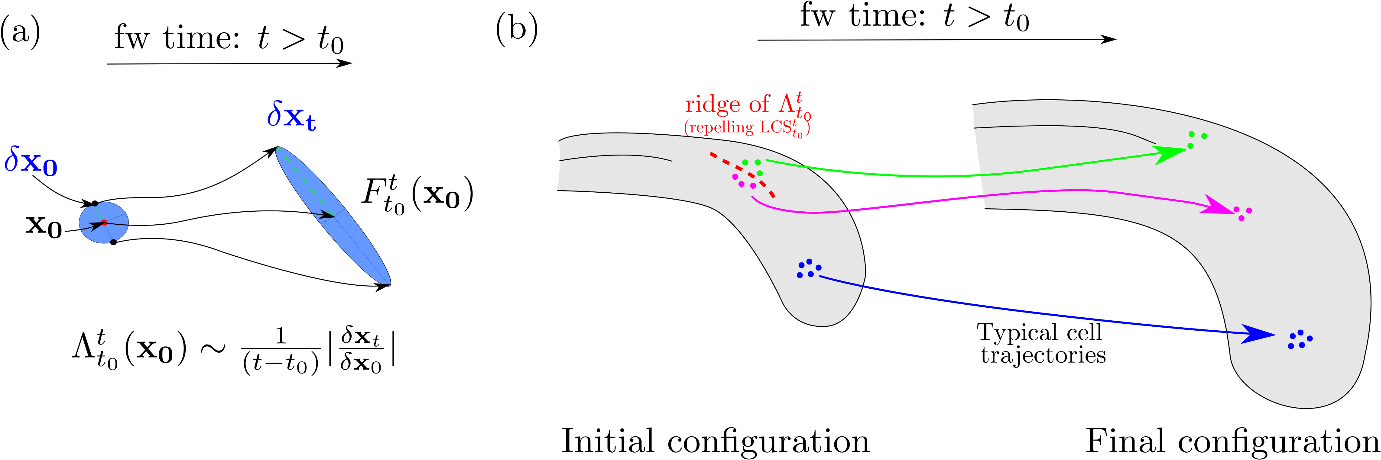
(a) The Lagrangian Stretching 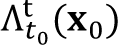 is a measure of the maximum separation generated by the flow over the time interval [*t_0_*, *t*] between two nearby points in the neighborhood of ***x***_0_. (b) Ridges (ie sets of high values) of 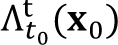 mark repelling LCS over [*t_0_*, *t*], which are the set of initial cell locations ***x***_0_ where the Lagrangian stretching is maximum.

